# Neuropathologically-directed profiling of *PRNP* somatic and germline variants in sporadic human prion disease

**DOI:** 10.1101/2024.06.25.600668

**Authors:** Gannon A. McDonough, Yuchen Cheng, Katherine Morillo, Ryan N. Doan, Connor J. Kenny, Aaron Foutz, Chae Kim, Mark L. Cohen, Brian S. Appleby, Christopher A. Walsh, Jiri G. Safar, August Yue Huang, Michael B. Miller

**Author notes:** Correspondence and requests for materials should be addressed to senior authors M.B.M. and A.Y.H. These authors contributed equally (Gannon A. McDonough, Yuchen Cheng).

## Abstract

Creutzfeldt-Jakob Disease (CJD), the most common human prion disease, is associated with pathologic misfolding of the prion protein (PrP), encoded by the *PRNP* gene. Of human prion disease cases, ∼1% were transmitted by misfolded PrP, ∼15% are inherited, and ∼85% are sporadic (sCJD). While familial cases are inherited through germline mutations in *PRNP*, the cause of sCJD is unknown. Somatic mutations have been hypothesized as a cause of sCJD, and recent studies have revealed that somatic mutations accumulate in neurons during aging. To investigate the hypothesis that somatic mutations in *PRNP* may underlie sCJD, we performed deep DNA sequencing of *PRNP* in 205 sCJD cases and 170 age-matched non-disease controls. We included 5 cases of Heidenhain variant sporadic CJD (H-sCJD), where visual symptomatology and neuropathology implicate focal initiation of prion formation, and examined multiple regions across the brain including in the affected occipital cortex. We employed Multiple Independent Primer PCR Sequencing (MIPP-Seq) with a median depth of >5,000X across the *PRNP* coding region and analyzed for variants using MosaicHunter. An allele mixing experiment showed positive detection of variants in bulk DNA at a variant allele fraction (VAF) as low as 0.2%. We observed multiple polymorphic germline variants among individuals in our cohort. However, we did not identify bona fide somatic variants in sCJD, including across multiple affected regions in H-sCJD, nor in control individuals. Beyond our stringent variant-identification pipeline, we also analyzed VAFs from raw sequencing data, and observed no evidence of prion disease enrichment for the known germline pathogenic variants P102L, D178N, and E200K. The lack of *PRNP* pathogenic somatic mutations in H-sCJD or the broader cohort of sCJD suggests that clonal somatic mutations may not play a major role in sporadic prion disease. With H-sCJD representing a focal presentation of neurodegeneration, this serves as a test of the potential role of clonal somatic mutations in genes known to cause familial neurodegeneration.

## Introduction

Prion diseases are transmissible, progressive, fatal neurodegenerative disorders associated with the pathologic misfolding and aggregation of the prion protein (PrP). Encoded by the *PRNP* gene, the normal cellular prion protein PrP^C^ serves as the substrate for the disease-specific conformer PrP^Sc^, the primary component of infectious prions. Prions can be transmitted between mammals, as entities such as scrapie in sheep and goats, chronic wasting disease in deer and elk, bovine spongiform encephalopathy in cows [13], and kuru in humans [26]. While prion diseases are distinct in their infectivity, other etiologies drive significant disease burden.

Prion disease affects 1-2 people per million annually, with a lifetime individual risk of 1 in 5,000 [63], with most cases designated as Creutzfeldt-Jakob disease (CJD) [83]. Human prion disease can arise from infectious, familial, or sporadic origins. Infectious transmission, including kuru, iatrogenically acquired disease, and zoonotic transmission of bovine spongiform encephalopathy, have caused <1% of cases [88]. Most prion disease cases (∼85%) are sporadic (sCJD), while ∼15% of cases are familial. Human prion disease presentations show a trimodal age of onset by etiology, with infectious/acquired cases peaking around age 30, genetic/inherited cases around age 50, and sporadic cases above age 60 [3]. The cause of sporadic prion disease is not known, but familial cases may offer insights into its pathogenesis.

Familial prion diseases result from inherited autosomal dominant gain-of-function germline mutations in the *PRNP* gene. Three fully-penetrant missense variants cause defined clinicopathologic syndromes: P102L causes Gerstmann-Straussler-Scheinker (GSS) syndrome [32–33], D178N causes fatal familial insomnia (FFI) and familial CJD (fCJD) [59], and E200K also causes fCJD [31, 63, 81]. In addition to these fully-penetrant missense variants, other variants, including V210I and V180I, confer increased risk [63]. Variants producing premature stop codons, when located near the N-terminus and thus truncating the majority of PrP, may be protective [63], fitting with the observation that deletion of *PRNP* in experimental animals prevents prion infection [11]. The missense variant G127V, discovered in kuru-exposed populations, appears protective against prion disease [58]. Codon 129 is a polymorphic locus that modulates prion disease pathogenesis and clinical presentation; for example, 129 methionine or valine in cis determines the neuropathological location and symptomatology of the D178N pathogenic variant allele [27, 44]. Genome-wide association studies (GWAS) have confirmed the *PRNP* locus as the dominant genetic influence on prion disease [56–57].

Unlike familial prion disease, the initiating mechanism of sporadic prion disease is not known. Since *PRNP* germline mutations can cause early-onset disease, we hypothesized that later-onset sporadic prion disease may result from somatic mutations, which arise not from parental inheritance but instead arise postzygotically during development and aging and are thus only present in a subset of cells within an individual. While somatic mutations are known as drivers of neoplasia [28, 81], they also develop in non-neoplastic tissues [29, 52], including in post-mitotic cells such as neurons where they accumulate with age and in neurodegeneration [50, 62].

In certain disease settings, somatic variants have resulted in later-onset or localized distribution of disease when compared to germline variants. For example, germline *TP53* mutations in Li-Fraumeni syndrome lead to earlier-onset cancers than somatic-only mutations [66]. In the brain, somatic mTOR pathway mutations in a wider distribution produce hemimegalencephaly [73–74], while more restricted mTOR pathway mutations produce focal cortical dysplasia [7, 19]. Even somatic mutations shared by a limited number of cells have the potential for biological significance [43]. In contrast with the early age range of familial prion disease, the later onset of sporadic prion disease suggests possible initiation by somatic mutations in *PRNP*.

Somatic mutations in familial disease genes present a compelling potential mechanism for sporadic later-onset neurodegenerative diseases in general. In Alzheimer’s disease and prion disease, previous studies have not demonstrated pathogenic somatic mutation as a common disease-causing mechanism [78, 84]. However, these studies focused on neuropathologically classic cases that proceed with widespread multifocal disease. The Heidenhain variant of sporadic Creutzfeldt-Jakob prion disease (H-sCJD) shows anatomically concentrated pathology in the occipital visual cortex [6, 15], offering the opportunity to evaluate potential pathogenic somatic mosaicism in focal neurodegenerative disease.

In the present study, we employ Multiple Independent Primer PCR Sequencing (MIPP-Seq) to perform high-coverage sequencing and sensitive somatic mutation detection on the *PRNP* gene in large cohorts, including H-sCJD, sCJD, and age-matched controls. For broad neuroanatomical profiling of H-sCJD, we sample a range of brain areas including multiple occipital cortex regions, to examine the pathogenicity of low-fraction somatic single-nucleotide variants (SNVs) and short insertions/deletions (indels) in *PRNP*.

## Materials and Methods

### Patient sample cohorts

Postmortem brain samples from two cohorts of individuals were used in this study: sCJD and non-disease controls. 205 cases of confirmed sCJD (**Fig. 2A, Supplemental Table 1**) were collected by the National Prion Disease Pathology Surveillance Center (NPDPSC) at Case Western Reserve University (Cleveland, OH), approved under IRB protocol 01-14-18 and 03-14-28. sCJD cases ranged from 45-86 years of age and were comprised of 95 females and 110 males, with a mean age at death of 68.2 years and a standard deviation of 8.8 years. CJD was confirmed using histopathology, immunohistochemistry, and Western blot analysis for PrP^Sc^, as previously described [24, 72], in combination with Sanger sequencing of *PRNP*, cerebrospinal fluid analysis, and patient clinical profiles. Familial prion disease cases were excluded from this study, based on Sanger sequencing for known germline *PRNP* disease-causing mutations, to validate the cohort as sCJD. The sCJD cohort included a subcohort of five cases of H-sCJD, defined as a pure visual clinical presentation with hypersensitivity in the occipital lobes on diffusion weighted imaging (DWI) [2]. Findings for sCJD in this study do not include H-sCJD, which is reported here as a separate category.

For non-disease controls, a cohort of 170 individuals, negative for any neurologic disease diagnosis and covering the age range of the sCJD cohort, was obtained from the NIH Neurobiobank (**Supplemental Table 1**). Control cases ranged from 35-89 years of age at death and were comprised of 55 females and 115 males, with a mean age of 61.0 years and a standard deviation of 12.7 years. The non-disease cohort was assembled by the NIH Neurobiobank from tissue obtained at the following repositories: the Human Brain and Spinal Fluid Resource Center of VA Greater Los Angeles Healthcare System in Los Angeles, CA; University of Miami Miller School of Medicine Brain Bank in Miami, FL; Mount Sinai Neuropathology Brain Bank and Research CoRE in New York, NY; University of Pittsburgh Neuropathology Brain Bank in Pittsburgh, PA; and the University of Maryland in Baltimore, Maryland. Postmortem tissues were collected according to their respective institutional protocols and examined in this study with the approval of the Boston Children’s Hospital Institutional Review Board (S07-02- 0087 with waiver of authorization, exempt category 4).

For all individuals (sCJD and non-disease controls), cerebellar cortex samples were selected for profiling, on the basis of sample availability and cerebellar involvement in many CJD cases. For H-sCJD, which exhibits pathology focused in the occipital cortex, a broader anatomic sampling included three occipital cortex areas (BA 17, 18, and 19), as well as prefrontal cortex (BA 46), parietal cortex, temporal cortex, thalamus, and the dentate nucleus of cerebellum (**Supplemental Table 1**). For a subset of the control individuals, additional brain regions were also profiled, for comparison with H-sCJD (**Fig. 1A**).

**Fig. 1.**
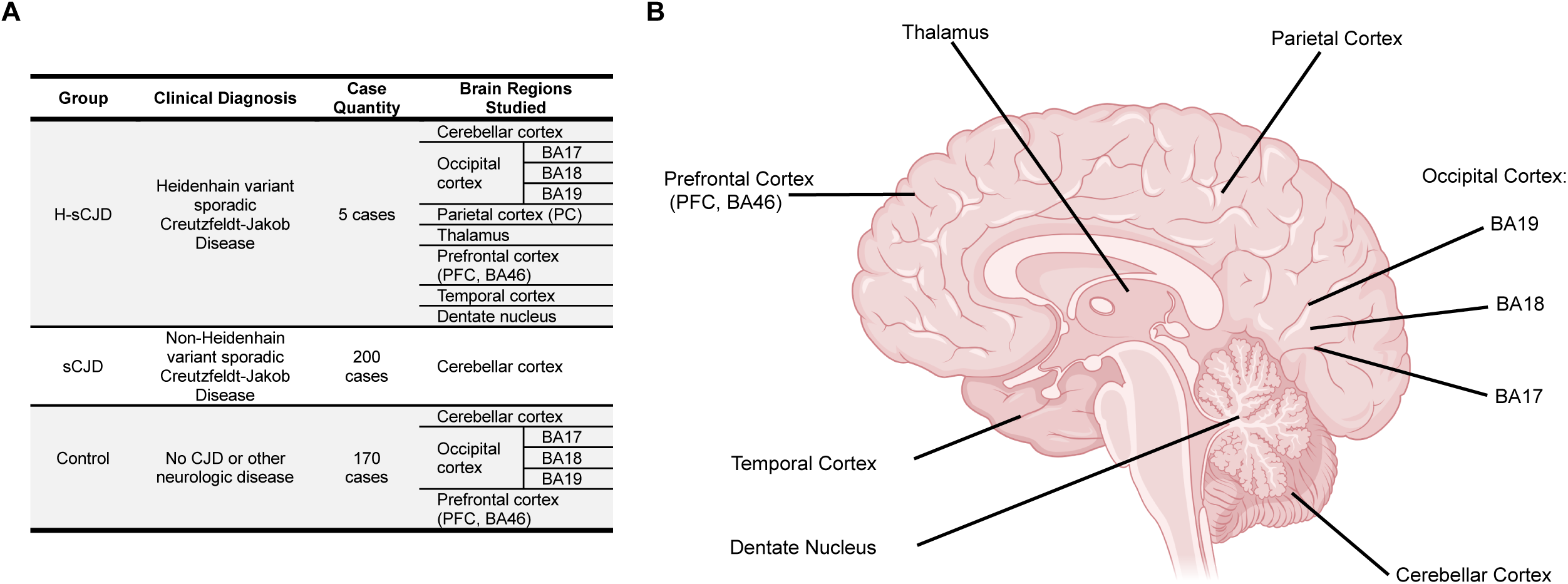
Study sample overview: cases and human brain regions. A.) Case summary. The study profiled cerebellar cortex tissue from 200 non-Heidenhain variant sporadic Creutzfeldt-Jakob Disease (sCJD) cases, and additional tissue from various regions in 5 Heidenhain variant sporadic Creutzfeldt-Jakob Disease (H-sCJD) cases, as well as from 170 non-disease control cases. B.) Brain regions studied for H-sCJD. Multiple areas of occipital cortex were profiled to broadly examine anatomic distribution of variants in H- sCJD, along with a diverse set of other brain regions, including parietal cortex, thalamus, prefrontal cortex, temporal cortex, and dentate nucleus of cerebellum. A set of control cases was also profiled with multiple occipital cortex sites and prefrontal cortex. In all sCJD and control cases, cerebellar cortex was studied.

**Fig. 2.**
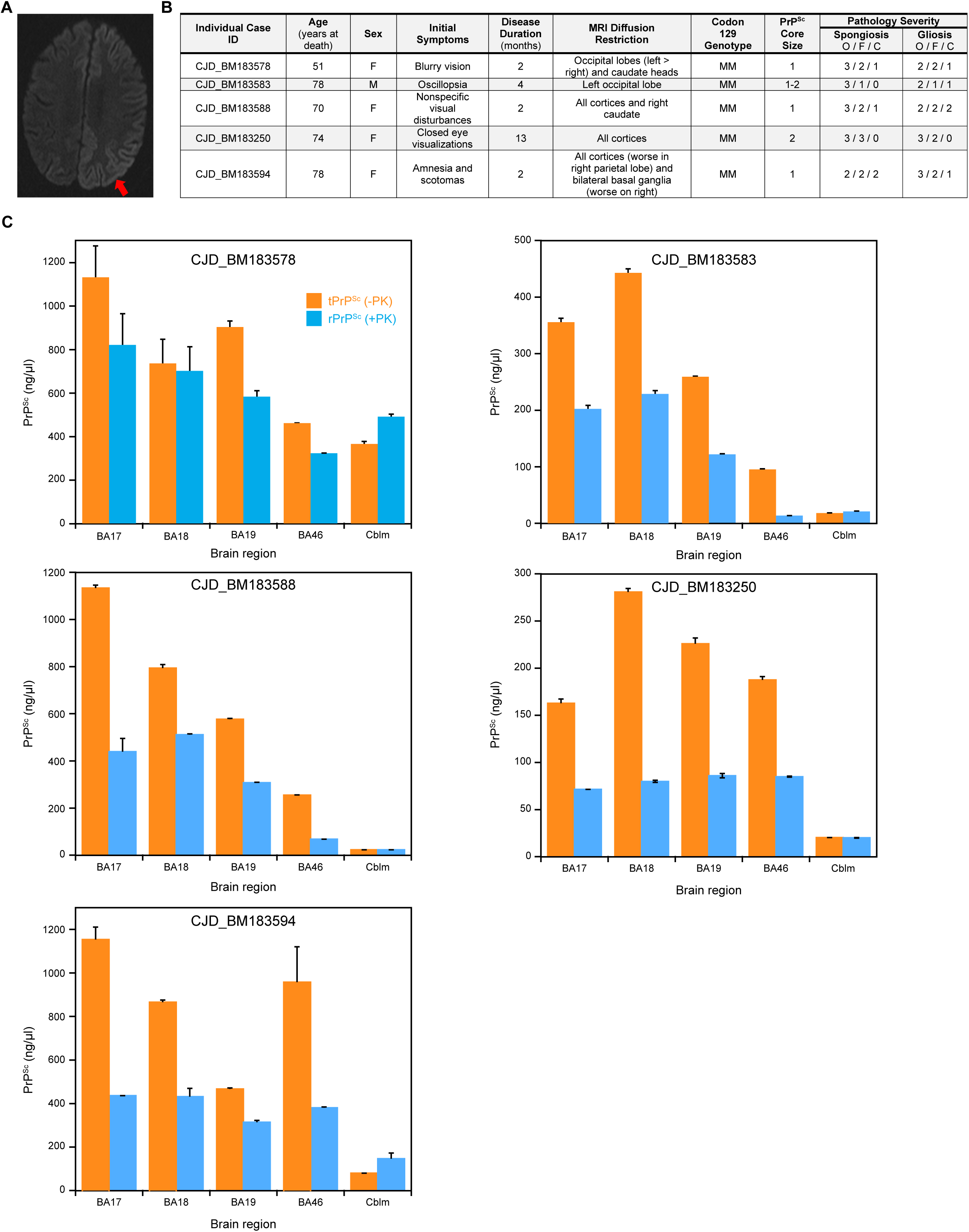
Heidenhain variant (H-sCJD) clinicopathologic and conformation-dependent immunoassay (CDI) findings. A.) Representative MRI of H-sCJD, axial view of hypersensitivity (arrow) in the cortical ribbon primarily affecting the left occipital and parietal cortices on diffusion weighted imaging. B.) Table summarizing H-sCJD clinicopathologic findings. PrP^Sc^ core size refers to the protease-resistant core size designation assayed by electrophoretic mobility on Western blot. Pathology severity is scored 0-3 for each case based on both spongiosis and gliosis findings, where a greater score indicates more pathology severity in a given brain region; O: occipital cortex, F: frontal cortex, C: cerebellum. C.) Conformation-dependent immunoassay (CDI) findings for each H-sCJD case. PrP^Sc^ abundance in ng/µL (y-axis) is measured before (orange) and after (blue) Proteinase K treatment. Examined brain regions are displayed on the x-axis; BA17, BA18, BA19: Occipital cortex- Brodmann area 17, 18, and 19, respectively, BA46: Prefrontal cortex- Brodmann area 46, and Cblm: cerebellum.

### Evaluation of Heindenhain-variant sporadic Creutzfeldt-Jakob Disease cases for total (tPrP^Sc^) and protease-resistant PrP^Sc^ (rPrP^Sc^) by Conformation-Dependent Immunoassay (CDI)

The Conformation-Dependent Immunoassay (CDI) was performed before and after Proteinase K treatment and after precipitation of PrP^Sc^ from brain homogenates with phosphotungstic acid as described previously [40, 42, 76, 77] with the following minor modifications. First, we used white Lumitrac 600 High Binding Plates (E&K Scientific, Santa Clara, California) coated with mAb 8H4 (PrP epitope 145-185) [87] in 200 mM NaH_2_PO_4_ containing 0.03% (w/v) NaN_3_, pH 7.5. Second, aliquots of 20 µL from each fraction containing 0.007% (v/v) of Patent Blue V (Sigma) were directly loaded into wells of white strip plates prefilled with 200 µL of Assay Buffer (Perkin Elmer, Waltham, Massachusetts). Finally, the captured PrP was detected by a europium-conjugated [77] anti-PrP mAb 3F4 (PrP epitope 108-112) and the time-resolved fluorescence (TRF) signals were measured by the multi-mode microplate reader PHERAstar Plus (BMG LabTech, Durham, North Carolina). The recHuPrP(90–231,129M) and PrP(23- 231,129V) used as calibrants in the CDI were prepared and purified as described previously [40, 42, 77].

### Bulk DNA extraction

Genomic DNA was extracted from freshly frozen brain tissue samples (∼25 mg) using commercially available extraction kits, Qiagen EZ1 DNA Tissue Kit (953034) or Qiagen DNeasy Blood & Tissue Kit (69504). DNA extraction from CJD samples was performed in biosafety level 3 conditions and included 5 M guanidinium hydrochloride denaturation to inactivate prion infectivity [25, 35], permitting downstream studies to be performed in standard molecular biology laboratory conditions. Extracted bulk DNA was quantitated using the Thermo Fisher Quant-iT PicoGreen DNA Quantification for gDNA Kit (33120, Invitrogen).

### PRNP amplicon sequencing by MIPP-Seq

We performed MIPP-Seq, for its demonstrated ability to reach sufficient coverage to accurately and precisely detect low-fraction somatic variants as previously described [21]. MIPP-Seq involves two sequential PCR steps, generating amplicons independently covering the genomic region of interest, which in this study consisted of at least 3 independent amplicons covering the coding region of the *PRNP* gene. The first PCR step uses primers that hybridize to the genomic target– one of eleven amplicons covering the *PRNP* gene (**Supplemental Table 2**). Step 1 primers also include Illumina adapter sequences upstream (5’) of the *PRNP*-targeting sequence, with the P5 adapter linked to the forward primer and the P7 adapter linked to the reverse primer. Then, the second round of PCR produces dual-indexed PCR amplicons with sample-specific barcodes.

For step 1 PCR, eleven overlapping amplicons targeting the entire *PRNP* protein-coding region (chr20:4679867-4680628 for the entire *PRNP* protein-coding region, chr20:4666797-4682234 for the fully transcribed *PRNP* gene, including exon 1, intron 1, and exon 2) were designed with the National Library of Medicine Genome Data Viewer on the human GRCh37 reference genome to extract the *PRNP* genomic locus and flanking sequence (**Fig. 3A**). The UCSC Genome Browser was also utilized for visualization and annotation of the *PRNP* gene. Primers were designed using Primer3Plus. Primers were designed to a target T_m_ of 60°C, with a minimum T_m_ of 59°C and a maximum T_m_ of 62°C. To ensure that all primers were unique and of similar amplicon length, amplicons had a target length of 225-300 bp, with adjustments to avoid primers within repeat sequences, specifically the *PRNP* octapeptide repeat region (OPR) in this study. Each pair of primer sequences was reviewed by in-silico PCR to ensure the production of a unique amplicon in the human genome. The full PCR step 1 oligonucleotides included random 5-nucleotide sequences for diversity as well as Illumina adapters, for a 5’-3’ sequence of adapter-5N-primer. Step 1 PCR (*PRNP*- targeting) was performed using 16 cycles with 60°C annealing temperature in a 25 μL reaction mix containing 50 ng of input isolated DNA, Phusion Hot Start II DNA Polymerase, dNTPs (10 mM each), Phusion HF buffer, and the forward and reverse primers (ThermoFisher, 0.5 μM each). The 11 PCR reactions for each DNA sample, corresponding to each of the 11 amplicons, were pooled and cleaned up with Polyethylene Glycol-8000 and Sera-Mag Speedbeads magnetic beads (Fisher 09-981- 123) [75], at 1.5:1 ratio by volume.

**Fig. 3.**
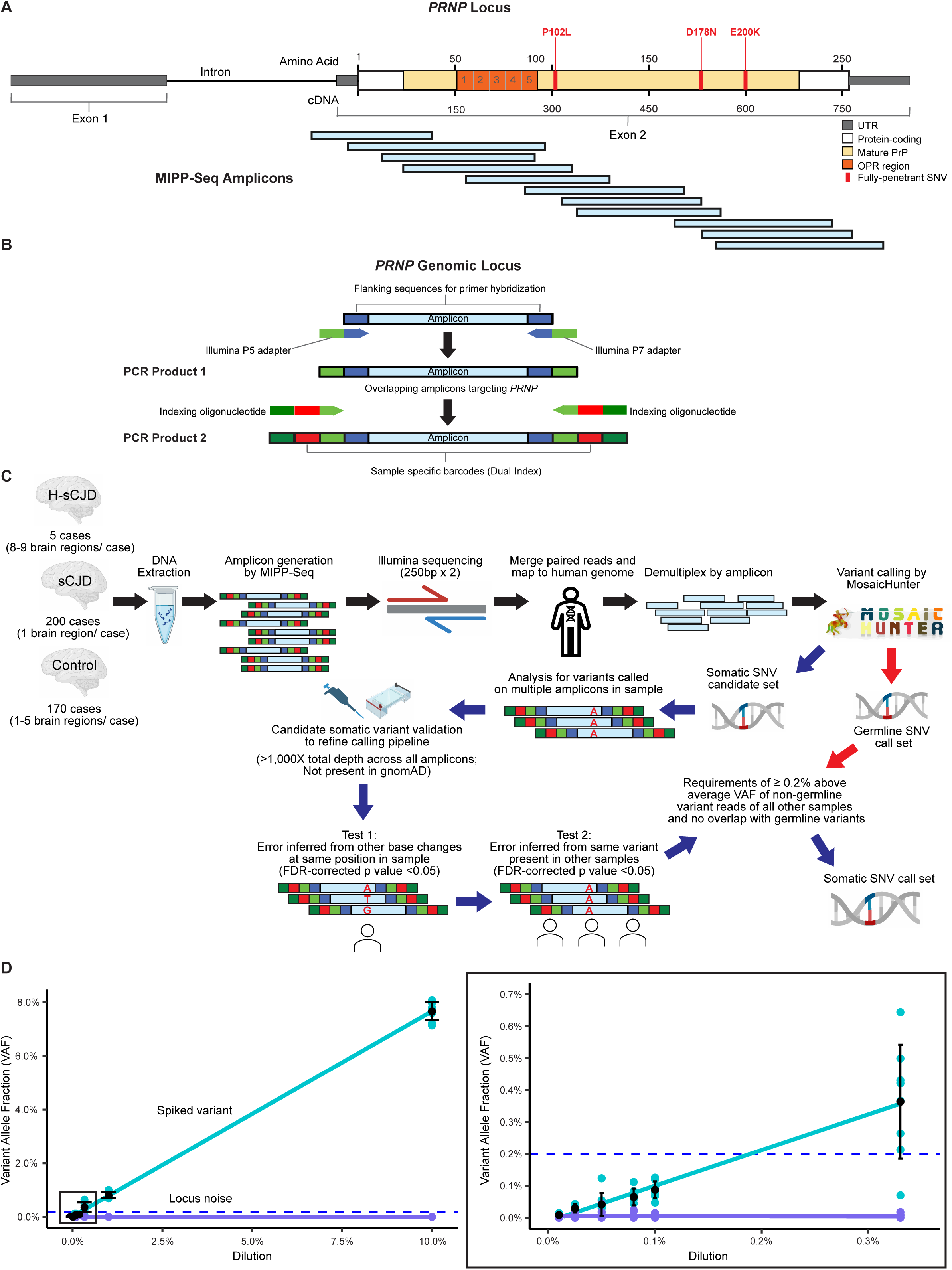
Approach and analysis for targeted deep sequencing of *PRNP gene*. A.) *PRNP* overlapping amplicon sequencing method. The *PRNP* coding region was targeted using Multiple Independent Primer PCR Sequencing (MIPP-Seq), which produced 11 overlapping amplicons ranging from 160 bp to 302 bp in length, followed by dual indexing PCR. B.) MIPP-Seq workflow. First, using primers hybridizing to the target region of *PRNP*, samples underwent PCR to produce amplicons including Illumina adapters. A second round of PCR was performed with indexing oligonucleotide primers to produce amplicons with dual-index sample-specific barcodes. C.) Overview of ultradeep targeted sequencing of *PRNP* coding region. First, DNA was extracted from Heidenhain and non-Heidenhain sCJD and control cases, followed by PCR with overlapping amplicons, dual indexing by PCR, and Illumina sequencing. Paired 250 bp reads were merged and mapped to the human genome (GRCh37) and demultiplexed by amplicon. Variant calling was performed by MosaicHunter and divided into two pathways: candidate somatic variants and germline variants. Following candidate somatic variant validation, we refined our pipeline to exclude candidate somatic variants that (1) did not possess at least 1,000X total depth across all covered amplicons and/or (2) were present in gnomAD, (3) overlapped with previously called germline variants, and/or (4) possessed a variant allele fraction (VAF) less than 0.2% above the average VAF of non-germline variant reads of all other samples. D.) M129V polymorphic allele mixing experiment. Using dilutions of bulk human DNA containing the known p.129V polymorphism and p.129M reference allele, we detected spiked variants above noise at that locus, with the blue dashed line indicating the 0.2% VAF threshold used for subsequent experiments.

Step 2 PCR was performed with oligonucleotides (ThermoFisher) to incorporate 8-nucleotide dual indexes to barcode all library products from each DNA sample. Forward primers included, from 5’ to 3’, 25-nucleotide IS5 indexing adapter, 8- nucleotide index, and 26-nucleotide sequence to hybridize with the P5 Illumina adapter incorporated in step 1 PCR [60]. Reverse primers included, from 5’ to 3’, 24-nucleotide IS6 indexing adapter, 8 nucleotide index, and 21 nucleotide P7 sequence to hybridize with the P7 Illumina adapter incorporated in step 1 PCR (**Fig. 3B**). Step 2 PCR reactions were performed in 25 μL volume containing purified Step 1 PCR product as input, indexing primers, Phusion Hot-Start II DNA Polymerase, dNTPs (10 mM each), and Phusion HF buffer. Step 2 PCR included an additional 14 cycles of amplification (60°C annealing temperature). Products were quantified (Quant-iT PicoGreen dsDNA Assay Kit, ThermoFisher), pooled, and doubly size-selected by 0.4:1 ratio exclusion and 0.8:1 ratio inclusion for 300-500bp library size fragments (AMPure XP magnetic beads, Beckman Coulter). These pooled libraries underwent 2 × 250 bp sequencing (Illumina HiSeq 2500, Psomagen, Rockville, MD).

### Detection of PRNP somatic variants with MosaicHunter

Raw paired reads from MIPP-Seq were first merged by USEARCH [22] and aligned to the human GRCh37 reference genome by BWA-MEM [46], after extracting and trimming 5-nucleotide unique molecular identifier (UMI) sequence from both ends. We then demultiplexed the aligned reads by amplicon, considering only reads that had 95% reciprocal overlap with the designed amplicon region and had four or less mismatches. PCR duplicate reads were removed by using the UMI and alignment information. We calculated the average deduplicated depth across amplicons for each sample and performed down-sampling to ensure the depth distribution is comparable between sCJD and control groups (**Supplemental Fig. 1B**).

For each amplicon, germline and somatic SNVs were called by MosaicHunter [34], by only considering reads with ≥20 alignment and base-calling quality at any genomic position that were outside of amplicon primer regions. We then integrated sequencing information from multiple amplicons of a given sample, and considered only variants that were called in ≥2 amplicons and had a total depth of >100X across all the covered amplicons. Candidate variants with VAF >80% and VAF between 30-80% were considered as homozygous and heterozygous germline variants, respectively, and all the remaining candidates were considered as potential somatic variants. To further remove potential technical artifacts for somatic variant calling, we only included candidate somatic variants that met criteria as follows (**Fig. 3C**). First, candidate somatic loci must have a total depth of >1,000X across all covered amplicons. Second, the read fraction of mutant allele across amplicons must be significantly higher than the fractions of two other non-reference alleles at the same position in the sample (FDR-corrected Wilcoxon Rank-Sum Test’s p-value <0.05). Third, to avoid artifactual detection of variants caused by errors of amplification or sequencing, the read fraction of mutant allele across amplicons in the sample must be significantly higher than the fraction in other non-germline samples (FDR-corrected Wilcoxon Rank-Sum Test’s p-value <0.05 and the VAF difference >0.2%). Fourth, to further avoid artifactual detection of variants caused by read misalignment or errors of amplification or sequencing (such as index hopping), the mutant allele must not be present in the human genome aggregation database (gnomAD) (v2.1.1 used for informatic comparison with GRCh37 human genome alignment in this study, with validation in gnomAD v4.0.0) [12, 36], an approach to artifact minimization also employed by other recent studies [1, 5, 47].

Apart from the above criteria of our informatic variant-calling pipeline, we reviewed the raw VAFs of various loci to evaluate for potential somatic variants below the limit of detection based on pipeline criteria (**Fig. 4-5**, **Supplemental Fig. 2-3**). Further, using the Integrated Genomics Viewer (IGV), we manually inspected the sequencing reads around specific loci (20:468070, 20:4680112, and 20:4680118) that had previously been reported as somatic variants [65], where the proximity to the OPR region raised concern for read misalignment. We determined that misalignment of reads from alleles with one OPR deletion (24 bp deletion) is responsible for certain candidate variant calls, rather than bona fide somatic variants (**Supplemental Fig. 2B-C**). While no somatic variants at these loci passed our calling pipeline, this manual inspection was relevant in the examination of raw reads at these loci.

**Fig. 4.**
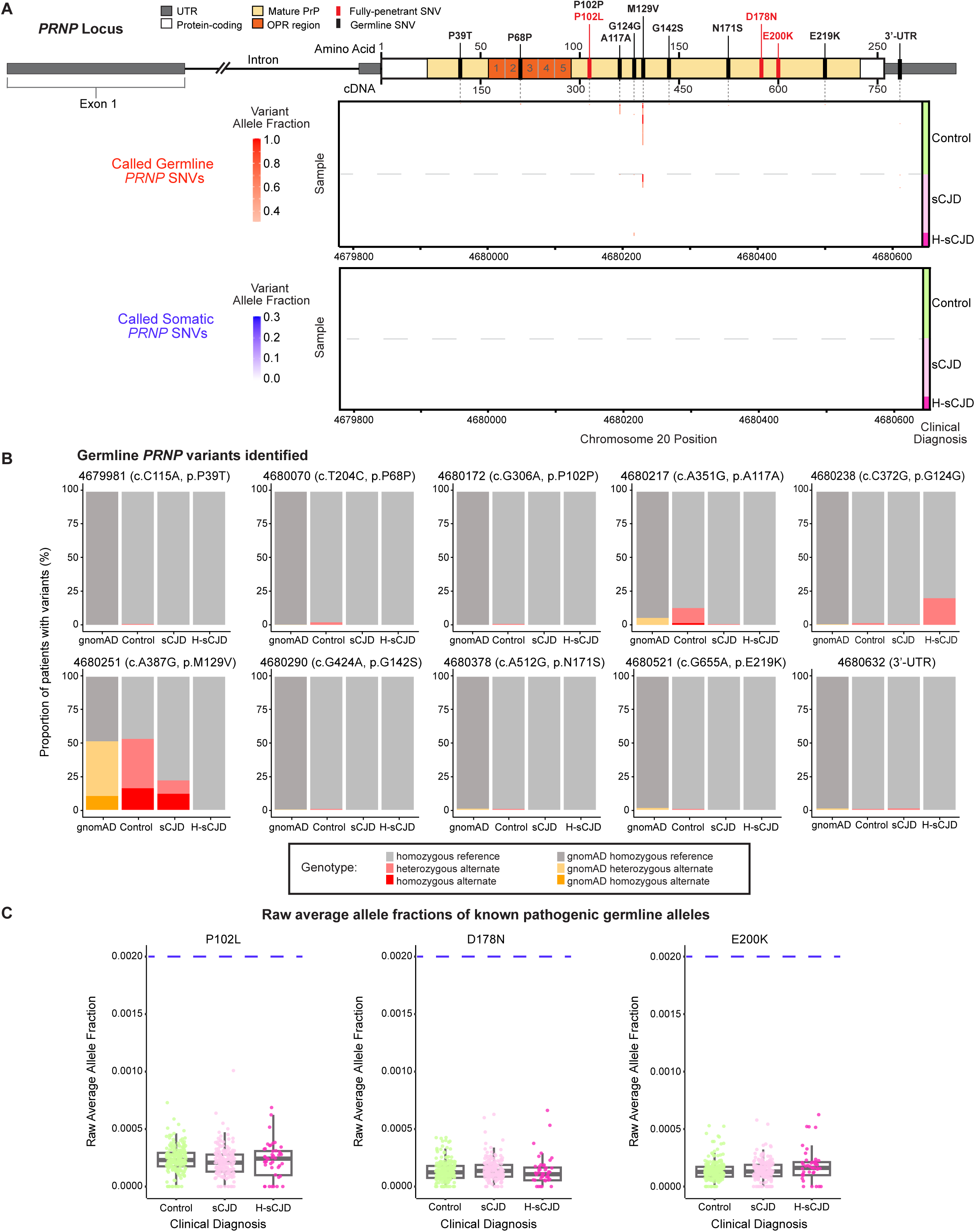
Somatic and germline single nucleotide variants (SNVs) in *PRNP* gene. A.) *PRNP* gene diagram with germline variants labeled and detected variant heatmaps showing chromosome position and samples. Within the gene diagram, the untranslated regions are represented in gray, the mature PrP region is represented in yellow including the octapeptide repeat region (OPR) is represented in orange, with the 5 repeats labeled 1-5. The amino acid coordinates are shown above and the cDNA coordinates are shown below the diagram. Fully penetrant familial *PRNP* variants (P102L, D178N, E200K) are labeled in red, and detected germline variants are identified in black, with corresponding location alignment within the germline variant heatmap (upper panel) and somatic variant heatmap (lower panel). Within each heatmap, covering the protein-coding region of *PRNP* in exon 2, non-disease control cases are displayed in the upper half, whereas sCJD cases are displayed in the lower half, with H-sCJD cases displayed at the bottom of each heatmap. Darker shading within the heatmaps represents a greater variant allele fraction (VAF) within a given sample at a variant locus. The germline (upper) heatmap contains detected variants ranging from 0.3-1.0 VAF, while the somatic (lower) heatmap contains detected variants ranging from 0.0-0.3 VAF. B.) Germline *PRNP* variants detected in this study. Bar plots depict the individual frequency of each genotype within each clinical diagnosis at each of the 10 detected germline variant loci, as well as the population frequency of this variant as described in gnomAD. Within our cohorts, light gray represents the homozygous reference genotype at that locus, light red represents the heterozygous alternate genotype, and dark red represents the homozygous alternate genotype. Within gnomAD, dark gray represents the homozygous reference genotype, light yellow represents the heterozygous alternate genotype, and dark yellow represents the homozygous alternate genotype. C.) Raw average allele fractions of the known pathogenic germline alleles. *PRNP* amplicon raw reads were analyzed for single nucleotide variants, upon removal of stringent variant calling and filtering that were implemented for standard analysis. Each clinical diagnosis cohort is shown with control samples in green, non-Heidenhain sCJD samples (sCJD) in light pink and Heidenhain sCJD (H-sCJD) samples in dark pink. Dashed blue lines show the 0.2% VAF detection threshold for the variant calling pipeline. Each data point represents a different sample from each individual case. Bonferroni-corrected Wilcoxon p-values for these plots showed no significance.

**Fig. 5.**
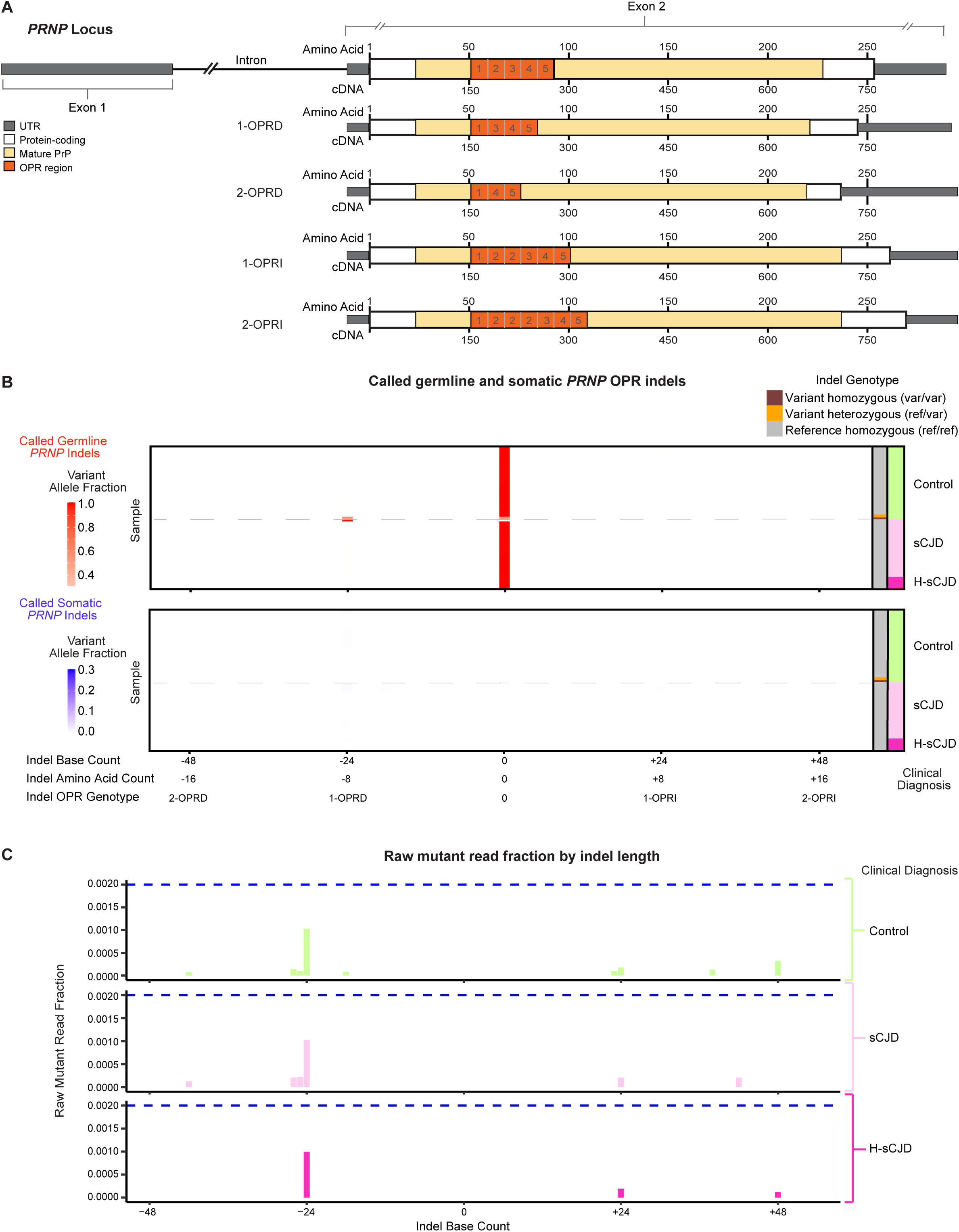
Somatic and germline octapeptide repeat region insertions and deletions (indels). A.) The *PRNP* reference locus is shown followed by diagrams of alterated OPR length. For example, 1-OPRD displays a deletion of one octapeptide repeat (loss of 8 consecutive amino acids, 24 nucleotides), while 2-OPRD displays a deletion of two octapeptide repeats (loss of 16 consecutive amino acids, 48 nucleotides). Conversely, 1-OPRI displays an insertion of one octapeptide repeat (gain of 8 consecutive amino acids, 24 nucleotides), while 2-OPRI displays an insertion of two octapeptide repeats (gain of 16 consecutive amino acids, 48 nucleotides). B.) Detected germline (upper panel) and somatic (lower panel) indels in OPR. Indel base count is displayed on the x-axes, with 0 representing the reference OPR, as well as the other indel genotypes indicated in part A. Non-disease control cases are displayed in the upper part of each heatmap, whereas sCJD cases are displayed in the lower part, with H-sCJD cases displayed at the bottom. Darker shading within the heatmaps represents greater VAF. As with the heatmaps in Figure 4, the upper germline heatmap contains detected variants ranging from 0.30-1.0 VAF, while the lower somatic heatmap contains detected variants ranging from 0.0-0.3 VAF. Samples are further categorized based on their germline OPR genotype at a given locus, homozygous for the variant genotype in brown, heterozygous genotype in orange, and homozygous for the reference allele in gray. C.) Raw mutant read fraction of each OPR indel type. *PRNP* amplicon raw reads were analyzed for OPR variants, upon removal of stringent variant calling and filtering that were implemented for standard analysis. Genomic loci spanning two octapeptide deletions (2-OPRD) to two octapeptide insertions (2-OPRI) are displayed on the x-axes and allele fraction is displayed on the y-axes. Each bar represents the raw mutant read fraction observed in the aggregate data of each sample cohort: controls, non- Heidenhain sCJD, and H-sCJD. Dashed blue lines show the 0.2% VAF detection threshold for the variant calling pipeline.

In addition to SNVs, we also developed a pipeline for calling somatic and germline insertions and deletions (indels) of the OPR of the *PRNP* gene. As amplicon PRNP 6 spans the full OPR region from both paired ends, we only considered reads derived from this amplicon for indel calling, to avoid artifacts introduced by read misalignment. The number of reads supporting the reference allele or any size of indel were calculated based on the CIGAR information of each read. The same depth and VAF cutoff used for SNV calling were applied to call germline and somatic indels of the OPR.

The targeted sequencing data generated in this study will be made available for research purposes limited to prion diseases (CJD, etc.), as specified by consent documents of enrolled participants.

### Sensitivity validation by 129 M/V allele mixing experiment

To establish somatic variant calling sensitivity, we performed a serial dilution of a known polymorphism, (chr20: 4680251, *PRNP* c.A385G, p.M129V). p.M129V is a common polymorphism in *PRNP*, with 69% M and 31% V alleles in the general population based on gnomAD, such that M/M and V/V homozygotes were sufficiently abundant for availability in this sensitivity experiment [44]. Extracted bulk DNA from a human tissue sample known to have 129V/V genotype was mixed into bulk DNA from a sample known to have 129M/M genotype, to produce a range of 129V variant allele fractions (VAFs; 10%, 1%, 0.33%, 0.1%, 0.08%, 0.05%, 0.025%, 0.01%). These mixtures each underwent MIPP-Seq and were analyzed using MosaicHunter as described. We evaluated for the lowest VAF at which the dilutant allele could be confidently distinguished from background noise, to determine the minimum VAF threshold for calling somatic variants, as depicted in **Fig. 3D**.

### Validation of potential somatic calls at selected PRNP loci to train variant-calling pipeline

To assist in establishing somatic variant calling parameters, we performed independent targeted-sequencing validation on three variant loci that were potential candidates with a preliminary calling threshold. For this, 150 bp of sequence upstream and downstream of each candidate locus was extracted to create a 300 bp sequence as the input for primer design. Primers were designed in Primer3Plus with the following parameters: 150 bp minimum and optimum amplicon length, 300 bp maximum amplicon length, 9 maximum 3’ stability, 12 maximum mispriming, 24 maximum pair mispriming, 18 bp minimum primer size, 20 bp optimum primer size, 27 bp maximum primer size, 55°C minimum primer T_m_, 60°C optimum primer T_m_, 65°C maximum primer T_m_, 5°C maximum T_m_ difference, 40% minimum primer GC%, 60% maximum primer GC%, 8 maximum self-complementarity, 3 maximum 3’ self-complementarity, 0 maximum #N’s, 5 maximum poly-X, 0 outside target penalty, 50 nM salt concentration, and 50 nM annealing oligo concentration.

Primers were designed to limit overlap with primers previously used in MIPP- Seq, and to ensure that each locus was covered by two different validation amplicons. Three amplicons were produced to cover these three loci, amplicon 007-343 (347 bp) covering the first two loci (forward: AACCTTGGCTGCTGGATG, T_m_ 59.8°C; reverse: CACCAGCCATGTGCTTCAT, T_m_ 60.7°C), amplicon 130-401 (272 bp) covering all three loci (forward: CCTGGAGGCAACCGCTAC, T_m_ 61.8°C; reverse: ATGGCACTTCCCAGCATGTA, T_m_ 61.5°C), and amplicon 348-597 (250 bp) covering the last locus (forward: AGCAGCTGGGGCAGTGGT, T_m_ 64.4°C; reverse: GGTGAAGTTCTCCCCCTTGG, T_m_ 63.1°C). Primer oligonucleotides (25 nM) were obtained from Genewiz. Targeted regions were amplified by PCR containing 50 ng input DNA, dNTPs (10 mM), 0.5 μM forward primer, 0.5 μM reverse primer, Phusion Hot-Start II DNA Polymerase, and Phusion HF buffer. PCR was performed as follows: initial denaturation stage at 98°C for 30 sec. (stage 1), followed by five annealing cycles beginning at 98°C for 10 sec., decreasing 1°C/ cycle until reaching 68°C, followed by 72°C for 30 sec. (stage 2), 28 extension cycles beginning at 98°C for 10 sec., 63°C for 30 sec., and 72°C for 30 sec. (stage 3), and finally 72°C for 10 min. (stage 4). PCR products were assessed for presence and specificity of target amplicons by 2% agarose gel electrophoresis. PCR products were purified using 2.3X AMPure XP beads. PCR products of different loci were pooled and sequenced (Illumina NovaSeq 2 × 250 bp, under Genewiz Amplicon-EZ service).

### Bioinformatic analysis of validation sequencing

Paired-end reads were merged by USEARCH [22], aligned to the human GRCh37 reference genome by BWA-MEM [46], and processed by GATK [18] for indel realignment. The aligned reads were then demultiplexed by the amplicon of each candidate and each sample, and the number of reads supporting each of the four nucleotides were counted. A mutation was considered validated if reads supporting the predicted mutant allele were three-fold more abundant than reads supporting the other two non-reference alleles.

## Results

### Creutzfeldt-Jakob disease and control human sample cohorts

To test the hypothesis that *PRNP* somatic mutations may cause sporadic Creutzfeldt-Jakob disease, we conducted amplicon-based targeted sequencing across the *PRNP* gene in extracted bulk DNA from human postmortem brain samples from sCJD and non-disease control cases. We examined three cohorts, designed to test specific potential biological impacts of *PRNP* somatic mutation (**Fig. 1A**).

The first cohort consisted of 5 individuals with H-sCJD, where visual symptoms and concentrated occipital neuropathology indicate an anatomically focal disease presentation, a pattern that is associated with causative somatic mutations in certain neurological conditions such as focal cortical dysplasia [20]. Therefore, H-sCJD represents a prion disease cohort that may have greater potential for etiologic contribution by somatic mutations in focal initiation of prion formation. In this cohort of clinically diagnosed cases, we confirmed the presence of occipital pathology by histological analysis for spongiosis and gliosis (**Fig. 2A**). We also performed the conformation-dependent immunoassay (CDI), which detected higher concentrations of protease-sensitive and protease-resistant PrP^Sc^ in the occipital cortex than the cerebellum in all five H-sCJD cases, with levels corresponding to those we published previously on a full phenotypic spectrum of sCJD [39–41]. Specifically, in three cases, the highest concentrations of PrP^Sc^ were detected in Brodmann area 17 (occipital primary visual cortex) with progressively lower levels in BA18 and BA19 (occipital visual association areas); whereas the other two showed the highest levels in BA18 (**Fig. 2B**). Frontal cortex, represented by BA46, showed variable levels of PrP^Sc^, while cerebellar cortex showed generally lower levels.

The second CJD cohort consisted of 200 individuals with non-Heidenhain sporadic CJD. The third cohort, serving as non-disease controls, consisted of 170 individuals with no known CJD or other neurologic disease. In all individuals, we studied cerebellar cortex, as a frequent site of pathology in CJD [53]. In H-sCJD cases, we examined additional brain regions, including occipital cortex regions (BA17, 18, and 19) for a detailed profiling of *PRNP* somatic mutation abundance in these specific areas implicated in disease. To provide a broad neuroanatomic survey in H-sCJD, we also examined parietal cortex, prefrontal cortex (BA46), temporal cortex, thalamus, and the dentate nucleus of the cerebellum (**Fig. 1B**). We also studied BA17, BA18, BA19, and BA46 in a subset of control cases, to allow for direct regional comparison with H-sCJD.

### Identification of SNVs in PRNP gene in human brain tissue

The *PRNP* gene plays a central role in prion disease risk and pathogenesis [48]. The *PRNP* locus is 16.2 kb in length and contains 2 exons and 1 intron. Exon 2 includes 762 bp encoding the 253 amino acids of full-length PrP (chr20: 4679867-4680628 in the human GRCh37 reference genome). PrP contains well-characterized structural and functional motifs, including the 123-bp octapeptide repeat region (OPR) (chr20: 4680017-4680139).

To evaluate somatic mutations in the *PRNP* gene, we utilized the MIPP-Seq method for high-fidelity targeted gene sequencing to identify low-fraction somatic variants in extracted bulk DNA from each sample (**Fig. 3A-C**). MIPP-Seq incorporates two consecutive PCR steps. In our application, the first PCR step used independent amplicons targeting overlapping areas of the *PRNP* protein-coding region, while the second PCR step generated dual-indexed libraries (**Fig. 3A-B**), which we then studied with 2 × 250bp Illumina DNA sequencing. We generated 11 overlapping amplicons targeting the *PRNP* protein-coding region for each case, and achieved a median depth of coverage of >5,000X in both prion disease and control cases, with 88% of all amplicons achieving at least 1,000X coverage (**Supplemental Fig. 1A-B**). Of specific relevance for the hypothesis of somatic *PRNP* mutation, this approach provided deep coverage of loci of fully-penetrant SNVs known to cause familial prion disease (**Supplemental Fig. 1C**).

Variants were called when detected on at least 2 amplicons with a total depth of >100X across all the covered amplicons. We performed an independent amplicon sequencing validation experiment on a preliminary set of candidate somatic variants, informing the construction of a somatic variant calling pipeline with high somatic variant calling stringency, requiring at least 0.2% greater VAF than the mean of the non-germline variant reads of all other samples (**Fig. 3C**). Previous studies on *PRNP* somatic mosaicism have highlighted the importance and challenges of sensitive and specific detection of low-fraction somatic variants in bulk tissue [45, 65, 86]. To assess the sensitivity of SNV calling, we performed a serial dilution of genomic DNA containing homozygous 129V/V polymorphic allele into genomic DNA containing homozygous 129M/M allele, producing a range of VAFs. We then performed MIPP-Seq and variant calling using MosaicHunter [34], which indicated that our method could confidently distinguish a true variant allele from background noise at a VAF as low as 0.2% or lower (**Fig. 3D**).

### Somatic and germline SNVs in PRNP in sCJD and control individuals

After performing MIPP-Seq across the *PRNP* protein-coding region for the set of 497 samples from 375 individuals from the three cohorts representing sCJD and controls, we identified SNVs. We first inspected for germline variants, determined by the VAF in our profiled brain tissue samples. We detected 10 distinct germline SNVs across the *PRNP* gene coding region and 3’ untranslated region (**Fig. 4A, upper panel**; **Fig. 4B**), most of which have been reported previously or are documented in the human genome aggregation database (gnomAD). As anticipated from established literature [14] and gnomAD [36], the V allele in the known polymorphic codon 129M/V was the most prevalent germline variant allele detected, and the relative abundance of this allele in our control cohort resembled that of gnomAD. We also observed an overrepresentation of homozygous reference 129M/M in sCJD, as has been previously reported [70]. Of note, in one control case, we detected a germline heterozygous SNV, p.P39T, which has not been previously reported or documented in gnomAD, though other variants at position 39 have been reported, including a likely incidental P39L variant in a case of frontotemporal dementia [68]. In H-sCJD cases, we detected only one germline SNV, p.G124G, a synonymous variant which was present but in a lower proportion of the control group and in the gnomAD database. The other germline variants that we detected were similarly abundant in control cases and in gnomAD, indicating that none of the germline variants detected in our study are pathogenic in prion disease.

Analysis for somatic SNVs in the *PRNP* gene produced no variants that were called by our sensitive but stringent pipeline, neither in the sCJD cohorts nor in the non-disease control group (**Fig. 4A, lower panel**). Notably, this includes specific sampling of all three main occipital cortex areas (BA17, 18, and 19) in the H-sCJD cases, a focal anatomic disease context where a somatic pathogenic *PRNP* variant might be suspected most. Correspondingly, we found no disease enrichment of somatic SNVs in sCJD cases as compared to controls. To examine for potential pathogenic somatic variants present at very low VAFs and thus potentially below the detection threshold of the stringent variant-calling pipeline, we analyzed the raw VAFs in each sample for known fully-penetrant germline *PRNP* variants (P102L, D178N, and E200K) [63]. We observed no significant somatic enrichment in these pathogenic variant alleles in prion disease cases over controls (**Fig. 4C**), and specific subset analyses found no somatic enrichment between different brain regions (**Supplemental Fig. 3A**) or between prion disease subtypes (**Supplemental Fig. 3B**).

### Somatic and germline OPR indels in PRNP in sCJD and control individuals

We next examined the length of the OPR, where certain germline variants of non-standard lengths are considered potentially pathogenic [55, 69] (**Fig. 5A**). The OPR consists of one nonapeptide followed by four consecutive octapeptide motifs, with the first two octapeptide units having identical reference sequences. The OPR is susceptible to expansions and deletions of one or more octapeptide units, and germline expansions of four or more octapeptide repeats potentially cause prion disease. We performed OPR sequence analysis using an amplicon (PRNP 6) where both reads span the full repeat region, with full-length OPR evaluation facilitated by the 2 x 250bp sequencing read length of this study.

We observed that all sCJD cases, including H-sCJD, possessed the reference OPR genotype with no germline octapeptide insertions or deletions, as expected based on the cohort design to exclude prion cases with known germline *PRNP* variants. A subset of control cases (5.9%, 10 individuals) possessed the germline heterozygous OPR genotype of a deletion of one octapeptide (1-OPRD), and 1 control individual (0.6%) was germline homozygous for 1-OPRD (**Fig. 5B, upper panel**). Given that these germline variants were detected exclusively in control cases, these appear to be benign polymorphisms, consistent with previous literature discussing the lack of pathogenicity of most OPR variants <4 octapeptides in length [30, 54, 71] and appearance of some such alleles in gnomAD.

Our examination for potential somatic octapeptide insertions or deletions, using our variant-calling pipeline, did not detect any somatic OPR indel variants in prion cases or controls (**Fig. 5B, lower panel**). Below the 0.2% VAF detection threshold of our stringent detection pipeline, we evaluated the raw OPR VAFs from each sample, where we only observed minimal reads across all the samples, with ∼0.1% mutant read fraction for 1-OPRD and <0.025% for other variants (**Fig. 5C**). These very low fractions of variant reads approximate intrinsic sequencing and amplification error rates in amplicon sequencing, and therefore do not appear to support a finding of somatic OPR variants in any of the samples.

## Discussion

While the origin of sporadic prion disease is not known, somatic mutation of the prion protein gene *PRNP* has been postulated as a potential cause. In this study, we profiled somatic mutations by performing deep (>5,000X) sequencing of the *PRNP* gene of multiple brain regions in the focal prion disease Heidenhain-variant sCJD, along with non-Heidenhain sCJD and non-disease controls. Using a custom MosaicHunter stringent somatic variant calling pipeline, we were able to identify somatic mutations at a VAF as low as 0.2%, which we demonstrated by an allele-mixing experiment. This builds on a recent *PRNP* somatic genetic study that reported 0.5% VAF sensitivity in a cohort of frontal lobe samples from 142 sCJD cases [65]. While our study focused on somatic variant detection, we incidentally cataloged 10 distinct germline SNVs in *PRNP* across our cohorts, including one which has not previously been reported, p.P39T, in a control individual. We also detected the germline deletion of one repeat within the OPR (1-OPRD) that is considered non-pathogenic, as it is present in unaffected individuals [8].

Our somatic mutation analysis did not identify any somatic SNVs or OPR indels — across H-sCJD, non-Heidenhain sCJD, and neurotypical controls — even with demonstrated sensitivity to 0.2% VAF. This stands in contrast to previous reports of various somatic *PRNP* variants in control and sCJD samples, primarily at 0.4-1.2% VAF in one study [65] and 1-7% VAF in a second study [86]. Apart from our variant-calling pipeline, we analyzed our raw sequencing reads for previously reported somatic SNVs and detected only low frequencies indicative of technical noise, all below 0.2% VAF, with no clear disease enrichment (**Supplemental Fig. 2**). Likewise, when examining raw VAFs for known germline pathogenic SNVs, we observed no disease enrichment, to a noise level below 0.1% VAF. Pipeline-independent examination of raw reads for OPR indels produced a similar finding, indicating that our method could reduce noise even below 0.2%, with no somatic variants detected. This application of repeat-spanning PCR amplification with longer length (2 x 250bp) Illumina sequencing of *PRNP* appeared to produce fewer artifactual OPR insertions/deletions than a recent Nanopore long-read sequencing approach [45].

We designed this study to investigate the role of somatic mutations in sporadic prion disease, with a focus on the Heidenhain variant, where focused occipital pathology suggested parallels with other focal neurologic disorders driven by somatic mutations. While focal cortical dysplasia, hemimegalencephaly, and mesial temporal lobe epilepsy are associated with clonal somatic variants detectable with bulk gene sequencing [19, 38, 73], we did not observe *PRNP* somatic mutations in the Heidenhain pathological focus in occipital cortex, nor in our survey of cerebellar tissue from 200 non-Heidenhain sCJD cases. With detection sensitivity of 0.2% VAF or lower, the lack of somatic mutations in *PRNP* in prion disease is striking, particularly in the affected occipital cortex in Heideinhain cases, and in the sampled cerebellum of the broader sCJD cohort. For investigation of the potential role of disease-causing somatic variants in neurodegeneration, focal prion disease would appear to be a highly suitable test case, given the outsized impact of causative germline genetics in prion disease (∼10- 15% of all cases) compared to other neurodegenerative diseases such as AD (<1%). Studies in AD have also not identified significant pathogenic brain somatic mutations in genes linked to causative germline disease, namely APP, PSEN1, and PSEN2 [37, 67, 78] apart from a single reported case with a 14% VAF variant in PSEN1 [9].

The relationship between pathogenic *PRNP* germline mutations and prion strain regional tropism carries potential implications for the possible role of somatic mutations in sporadic prion disease. As specific pathogenic mutations are associated with particular regional pathology distributions, the potential somatic occurrence of a known germline disease variant may be restricted to causing a similar pathology. By this logic, sporadic CJD would more likely be caused by somatic E200K, or by somatic D178N in cis with polymorphic 129V, which associate with a CJD neuropathological distribution. Similarly, sporadic fatal insomnia would be more likely linked with somatic D178N in cis with polymorphic 129M, which is associated with the thalamic-focused presentation of FFI [16]. Conversely, in this vein, somatic P102L would be less likely to cause sporadic prion disease, since non-familial presentations of GSS have not been reported. By this logic, given the unique neuropathological phenotype of some cases of sporadic prion disease, like H-sCJD that does not match a familial phenotype, a causative somatic mutation should be novel. Given the extensive range of germline *PRNP* variants detected in population genomic analyses, some families may carry such a hypothetical somatic variant unless it conferred embryonic lethality. This reasoning therefore would constrain the hypothesis of somatic *PRNP* mutations causing sporadic prion disease.

While the somatic mutation hypothesis has been considered an attractive potential initiating mechanism for sporadic prion disease, our negative findings align with projections from recent studies of human development on clonal somatic mutations, which are shared by multiple cells due to cellular proliferation after mutation incidence. Clonal somatic mutations of at least 0.2% variant allele fraction are relatively sparse [10] and are estimated to occur by around the 256-cell stage, or eighth cell division, of embryonic development (50% VAF x 0.5^8^ = 0.2% VAF). During the 255 divisions to reach the 256-cell stage, at a rate of ∼1.37 clonal somatic mutations per division [85], each individual would develop ∼349 detectable clonal somatic mutations. If these fall evenly across the 3.1 billion bases in the human genome [89], then each individual would have substantially fewer than one clonal somatic mutation (8.6 x 10^-5^) across the 762 bases in the *PRNP* coding sequence. Thus, the likelihood of a person having a clonal mutation in the *PRNP* gene is less than the lifetime risk of sporadic prion disease (1 in ∼5000) [63]. In this light, it is unsurprising that we did not detect such somatic mutations in our prion disease and control cohorts, indicating against clonal somatic mutation causality for sporadic prion disease.

The dearth of *PRNP* somatic mutations that we observe suggests that sporadic prion disease instead may be initiated by other mechanisms, such as spontaneous *de novo* misfolding of wild-type PrP [17, 82]. However, the lack of detected somatic variants in the brain, after neurodegenerative disease has progressed to death, does not fully disprove somatic variants as disease initiators. Given the capacity for very small amounts of misfolded PrP to propagate across the brain [61, 80], a single cell carrying a somatic pathogenic *PRNP* variant could be sufficient to trigger disease. Considering beyond clonal somatic mutations, which are shared by multiple cells, private mutations are those that are present in a single cell. Indeed, each neuron contains at least hundreds of private somatic mutations [49, 62], such that across the brain of a 65 year-old human, there would be nearly 10,000 neurons with each possible SNV (∼1000 private somatic SNVs per neuron [51] x ∼86 billion total neurons in human brain [4] = ∼86 trillion total SNVs in human brain ÷ 3.1 billion unique bases in genome of each human neuron [89] = ∼27,742 SNVs per base ÷ 3 possible substitutions at each base = ∼9,247 instances of each SNV in a human brain, which would essentially all fall in unique neurons). Of note, the figure of ∼9,247 of each SNV in each individual assumes equal chances of each SNV type, whereas C>T transitions at CpG sites (which are the cause of P102L, D178N, and E200K) are slightly more common than the mean somatic SNV in aging neurons [62], though private somatic CpG variants do not show the more pronounced enrichment seen in germline de novo mutations [79]. However, while a single cell could potentially initiate disease, private variants alone may be insufficient to cause prion disease in a typical human lifespan, as only a small proportion of the population develops disease, despite the estimated abundance of each somatic variant in the brain. If a single-neuron mutation is not enough to initiate sporadic prion disease, then investigations may more logically focus on clonal somatic mutations, which are shared by multiple cells of common lineage. Of note, a disease- originating mutated single cell or small clone may not survive until postmortem analysis, and indeed may be among the first cells to die, residing at the pathological epicenter [23].

There are several limitations in our study that should be considered to place the findings in proper context. In evaluating for somatic mutations that may be at very low VAFs or even lost due to cell death, tissue and cell sampling may not have captured the affected brain region or cells. This is of particular impact in a disease where a pathologic phenotype can be amplified by protein misfolding propagation from a small population of cells or even a single cell. However, in the H-sCJD cases in our study, the focus on occipital cortex and broad extra-occipital profiling significantly mitigates against the possibility of incomplete regional brain sampling. Our study is also limited to the somatic mutation VAF threshold of 0.2%, which may not detect ultra-low-fraction somatic mutations. Moreover, this study was performed with bulk genomic DNA and amplicon targeting, and thus is likely limited to detecting clonal somatic mutations but not private mutations unique to single cells. Targeted approaches using recently developed single-molecule sequencing [1] may be informative, especially for capturing ultra-low fraction somatic mutations, which may be further extended by single-cell approaches. Future studies may also benefit from isolation of particular cell populations identified by markers for neurons or glia, cell death, or for partially fragmented nuclei. Microdissection for particular local pathologic changes, including vacuolation density or PrP deposition, may also offer opportunities to pursue areas of greater somatic mutation potential.

Overall, our study using deep sequencing of the *PRNP* gene across 205 sCJD brains does not identify a significant role for *PRNP* somatic mutations in sporadic human prion disease. As focal prion disease represents a high-probability test for somatic mutation initiation of neurodegeneration, somatic variants in germline- implicated genes may not be a frequent cause of sporadic neurodegenerative disease. Knowledge of the likelihood of somatic *PRNP* variants causing sporadic prion disease also carries public health implications, as antisense oligonucleotide and other PrP-lowering strategies are investigated for patients with pathogenic mutations [64].

## Acknowledgements

Human tissue was obtained from the National Prion Disease Pathology Surveillance Center, the NIH Neurobiobank at the University of Maryland, the Human Brain and Spinal Fluid Resource Center of VA Greater Los Angeles Healthcare System, University of Miami Miller School of Medicine Brain Bank, Mount Sinai Neuropathology Brain Bank and Research CoRE, and University of Pittsburgh Neuropathology Brain Bank. We thank the donors and families for their contributions.

The National Prion Disease Pathology Surveillance Center was supported by J.G.S. CDC U51 CK000309 (to J.G.S.). J.G.S. was supported by NIH R01 NS074317, and R01 NS103848. M.B.M. was supported by NIH Director’s New Innovator Award DP2 AG086138, R01 AG082346, K08 AG065502, Doris Duke Foundation Clinical Scientist Development Award 2021183, T32 HL007627, the donors of the Alzheimer’s Disease Research program of the BrightFocus Foundation A20201292F, and the Brigham and Women’s Hospital Program for Interdisciplinary Neuroscience through a gift from L. and T. Rand. A.Y.H. was supported by R56 AG079857 and the Alzheimer’s Association Research Fellowship. C.A.W. is an Investigator of the Howard Hughes Medical Institute. Parts of Figures 1 and 3 were generated using BioRender. We thank Surachai Supattapone for helpful discussions.

## Contributions

M.B.M. conceived and designed the study, with input from C.A.W., J.G.S, and R.N.D. In discussion with M.B.M., J.G.S. selected and obtained prion disease brain specimens at the National Prion Pathology Surveillance Center, with input from B.S.A. on clinical phenotypes and brain MRI findings. M.L.C. performed neuropathological evaluation on prion disease tissue, and J.G.S. confirmed sCJD and H-sCJD diagnoses. C.K. and A.F. prepared prion disease tissue and performed bulk gDNA extraction. M.B.M. selected and obtained neurotypical control tissue. M.B.M. and K.M. extracted bulk gDNA from control samples. M.B.M. and K.M. performed MIPP-Seq in consultation with R.N.D., with assistance from C.J.K. M.B.M. and K.M. performed the M129V polymorphism mixing experiment. Y.C. and A.Y.H. performed computational analyses, including variant calling. G.A.M. and Y.C. designed and performed somatic variant validation experiments. M.B.M. and A.Y.H. supervised the study. G.A.M., M.B.M., A.Y.H., and Y.C. wrote the manuscript.

## References

1. Abascal F, Harvey LMR, Mitchell E, Lawson ARJ, Lensing SV, Ellis P, Russell AJC, Alcantara RE, Baez-Ortega A, Wang Y, Kwa EJ, Lee-Six H, Cagan A, Coorens THH, Chapman MS, Olafsson S, Leonard S, Jones D, Machado HE, Davies M, Øbro NF, Mahubani KT, Allinson K, Gerstung M, Saeb-Parsy K, Kent DG, Laurenti E, Stratton MR, Rahbari R, Campbell PJ, Osborne RJ, Martincorena I (2021) Somatic mutation landscapes at single-molecule resolution. Nature 593:405–410. doi: 10.1038/s41586-021-03477-4

2. Appleby BS, Appleby KK, Crain BJ, Onyike CU, Wallin MT, Rabins PV (2009) Characteristics of established and proposed sporadic Creutzfeldt-Jakob disease variants. Arch Neurol 66:208–215. doi: 10.1001/archneurol.2008.533

3. Appleby BS, Appleby KK, Rabins PV (2007) Does the presentation of Creutzfeldt-Jakob disease vary by age or presumed etiology? A meta-analysis of the past 10 years. J Neuropsychiatry Clin Neurosci 19:428–435. doi: 10.1176/jnp.2007.19.4.428

4. Azevedo FAC, Carvalho LRB, Grinberg LT, Farfel JM, Ferretti REL, Leite REP, Jacob Filho W, Lent R, Herculano-Houzel S (2009) Equal numbers of neuronal and nonneuronal cells make the human brain an isometrically scaled-up primate brain. J Comp Neurol 513:532–541. doi: 10.1002/cne.21974

5. Bae JH, Liu R, Roberts E, Nguyen E, Tabrizi S, Rhoades J, Blewett T, Xiong K, Gydush G, Shea D, An Z, Patel S, Cheng J, Sridhar S, Liu MH, Lassen E, Skytte A-B, Grońska-Pęski M, Shoag JE, Evrony GD, Parsons HA, Mayer EL, Makrigiorgos GM, Golub TR, Adalsteinsson VA (2023) Single duplex DNA sequencing with CODEC detects mutations with high sensitivity. Nat Genet 55:871–879. doi: 10.1038/s41588-023-01376-0

6. Baiardi S, Capellari S, Ladogana A, Strumia S, Santangelo M, Pocchiari M, Parchi P (2016) Revisiting the Heidenhain Variant of Creutzfeldt-Jakob Disease: Evidence for Prion Type Variability Influencing Clinical Course and Laboratory Findings. J Alzheimers Dis 50:465–476. doi: 10.3233/JAD-150668

7. Baldassari S, Ribierre T, Marsan E, Adle-Biassette H, Ferrand-Sorbets S, Bulteau C, Dorison N, Fohlen M, Polivka M, Weckhuysen S, Dorfmüller G, Chipaux M, Baulac S (2019) Dissecting the genetic basis of focal cortical dysplasia: a large cohort study. Acta Neuropathol 138:885–900. doi: 10.1007/s00401-019-02061-5

8. Beck JA, Poulter M, Campbell TA, Adamson G, Uphill JB, Guerreiro R, Jackson GS, Stevens JC, Manji H, Collinge J, Mead S (2010) PRNP allelic series from 19 years of prion protein gene sequencing at the MRC Prion Unit. Hum Mutat 31:E1551–63. doi: 10.1002/humu.21281

9. Beck JA, Poulter M, Campbell TA, Uphill JB, Adamson G, Geddes JF, Revesz T, Davis MB, Wood NW, Collinge J, Tabrizi SJ (2004) Somatic and germline mosaicism in sporadic early-onset Alzheimer’s disease. Hum Mol Genet 13:1219– 1224. doi: 10.1093/hmg/ddh134

10. Bizzotto S, Dou Y, Ganz J, Doan RN, Kwon M, Bohrson CL, Kim SN, Bae T, Abyzov A, NIMH Brain Somatic Mosaicism Network, Park PJ, Walsh CA (2021) Landmarks of human embryonic development inscribed in somatic mutations. Science 371:1249–1253. doi: 10.1126/science.abe1544

11. Büeler H, Aguzzi A, Sailer A, Greiner RA, Autenried P, Aguet M, Weissmann C (1993) Mice devoid of PrP are resistant to scrapie. Cell 73:1339–1347. doi: 10.1016/0092-8674(93)90360-3

12. Chen S, Francioli LC, Goodrich JK, Collins RL, Kanai M, Wang Q, Alföldi J, Watts NA, Vittal C, Gauthier LD, Poterba T, Wilson MW, Tarasova Y, Phu W, Grant R, Yohannes MT, Koenig Z, Farjoun Y, Banks E, Donnelly S, Gabriel S, Gupta N, Ferriera S, Tolonen C, Novod S, Bergelson L, Roazen D, Ruano-Rubio V, Covarrubias M, Llanwarne C, Petrillo N, Wade G, Jeandet T, Munshi R, Tibbetts K, Genome Aggregation Database Consortium, O’Donnell-Luria A, Solomonson M, Seed C, Martin AR, Talkowski ME, Rehm HL, Daly MJ, Tiao G, Neale BM, MacArthur DG, Karczewski KJ (2024) A genomic mutational constraint map using variation in 76,156 human genomes. Nature 625:92–100. doi: 10.1038/s41586-023-06045-0

13. Collinge J (2016) Mammalian prions and their wider relevance in neurodegenerative diseases. Nature 539:217–226. doi: 10.1038/nature20415

14. Collinge J, Palmer MS, Dryden AJ (1991) Genetic predisposition to iatrogenic Creutzfeldt-Jakob disease. Lancet 337:1441–1442. doi: 10.1016/0140-6736(91)93128-v

15. Cooper SA, Murray KL, Heath CA, Will RG, Knight RSG (2005) Isolated visual symptoms at onset in sporadic Creutzfeldt-Jakob disease: the clinical phenotype of the “Heidenhain variant.” Br J Ophthalmol 89:1341–1342. doi: 10.1136/bjo.2005.074856

16. Cracco L, Appleby BS, Gambetti P (2018) Fatal familial insomnia and sporadic fatal insomnia. Handb Clin Neurol 153:271–299. doi: 10.1016/B978-0-444-63945-5.00015-5

17. Deleault NR, Harris BT, Rees JR, Supattapone S (2007) Formation of native prions from minimal components in vitro. Proc Natl Acad Sci U S A 104:9741–9746. doi: 10.1073/pnas.0702662104

18. DePristo MA, Banks E, Poplin R, Garimella KV, Maguire JR, Hartl C, Philippakis AA, del Angel G, Rivas MA, Hanna M, McKenna A, Fennell TJ, Kernytsky AM, Sivachenko AY, Cibulskis K, Gabriel SB, Altshuler D, Daly MJ (2011) A framework for variation discovery and genotyping using next-generation DNA sequencing data. Nat Genet 43:491–498. doi: 10.1038/ng.806

19. D’Gama AM, Geng Y, Couto JA, Martin B, Boyle EA, LaCoursiere CM, Hossain A, Hatem NE, Barry BJ, Kwiatkowski DJ, Vinters HV, Barkovich AJ, Shendure J, Mathern GW, Walsh CA, Poduri A (2015) Mammalian target of rapamycin pathway mutations cause hemimegalencephaly and focal cortical dysplasia. Ann Neurol 77:720–725. doi: 10.1002/ana.24357

20. D’Gama AM, Woodworth MB, Hossain AA, Bizzotto S, Hatem NE, LaCoursiere CM, Najm I, Ying Z, Yang E, Barkovich AJ, Kwiatkowski DJ, Vinters HV, Madsen JR, Mathern GW, Blümcke I, Poduri A, Walsh CA (2017) Somatic Mutations Activating the mTOR Pathway in Dorsal Telencephalic Progenitors Cause a Continuum of Cortical Dysplasias. Cell Rep 21:3754–3766. doi: 10.1016/j.celrep.2017.11.106

21. Doan RN, Miller MB, Kim SN, Rodin RE, Ganz J, Bizzotto S, Morillo KS, Huang AY, Digumarthy R, Zemmel Z, Walsh CA (2021) MIPP-Seq: ultra-sensitive rapid detection and validation of low-frequency mosaic mutations. BMC Med Genomics 14:47. doi: 10.1186/s12920-021-00893-3

22. Edgar RC (2010) Search and clustering orders of magnitude faster than BLAST. Bioinformatics 26:2460–2461. doi: 10.1093/bioinformatics/btq461

23. Faucheux BA, Privat N, Brandel J-P, Sazdovitch V, Laplanche J-L, Maurage C-A, Hauw J-J, Haïk S (2009) Loss of cerebellar granule neurons is associated with punctate but not with large focal deposits of prion protein in Creutzfeldt-Jakob disease. J Neuropathol Exp Neurol 68:892–901. doi: 10.1097/NEN.0b013e3181af7f23

24. Foutz A, Appleby BS, Hamlin C, Liu X, Yang S, Cohen Y, Chen W, Blevins J, Fausett C, Wang H, Gambetti P, Zhang S, Hughson A, Tatsuoka C, Schonberger LB, Cohen ML, Caughey B, Safar JG (2017) Diagnostic and prognostic value of human prion detection in cerebrospinal fluid. Ann Neurol 81:79–92. doi: 10.1002/ana.24833

25. Frontzek K, Carta M, Losa M, Epskamp M, Meisl G, Anane A, Brandel J-P, Camenisch U, Castilla J, Haïk S, Knowles T, Lindner E, Lutterotti A, Minikel EV, Roiter I, Safar JG, Sanchez-Valle R, Žáková D, Hornemann S, Aguzzi A, THAUTAN-MC Study Group (2020) Autoantibodies against the prion protein in individuals with PRNP mutations. Neurology 95:e2028–e2037. doi: 10.1212/WNL.0000000000009183

26. Gajdusek DC, Zigas V (1959) Kuru; clinical, pathological and epidemiological study of an acute progressive degenerative disease of the central nervous system among natives of the Eastern Highlands of New Guinea. Am J Med 26:442–469. doi: 10.1016/0002-9343(59)90251-7

27. Gambetti P, Parchi P, Petersen RB, Chen SG, Lugaresi E (1995) Fatal familial insomnia and familial Creutzfeldt-Jakob disease: clinical, pathological and molecular features. Brain Pathol 5:43–51. doi: 10.1111/j.1750-3639.1995.tb00576.x

28. Garraway LA, Lander ES (2013) Lessons from the cancer genome. Cell 153:17–37. doi: 10.1016/j.cell.2013.03.002

29. Genovese G, Kähler AK, Handsaker RE, Lindberg J, Rose SA, Bakhoum SF, Chambert K, Mick E, Neale BM, Fromer M, Purcell SM, Svantesson O, Landén M, Höglund M, Lehmann S, Gabriel SB, Moran JL, Lander ES, Sullivan PF, Sklar P, Grönberg H, Hultman CM, McCarroll SA (2014) Clonal hematopoiesis and blood-cancer risk inferred from blood DNA sequence. N Engl J Med 371:2477–2487. doi: 10.1056/NEJMoa1409405

30. Goldfarb LG, Brown P, Little BW, Cervenáková L, Kenney K, Gibbs CJ Jr, Gajdusek DC (1993) A new (two-repeat) octapeptide coding insert mutation in Creutzfeldt-Jakob disease. Neurology 43:2392–2394. doi: 10.1212/wnl.43.11.2392

31. Goldgaber D, Goldfarb LG, Brown P, Asher DM, Brown WT, Lin S, Teener JW, Feinstone SM, Rubenstein R, Kascsak RJ (1989) Mutations in familial Creutzfeldt-Jakob disease and Gerstmann-Sträussler-Scheinker’s syndrome. Exp Neurol 106:204–206. doi: 10.1016/0014-4886(89)90095-2

32. Hsiao K, Baker HF, Crow TJ, Poulter M, Owen F, Terwilliger JD, Westaway D, Ott J, Prusiner SB (1989) Linkage of a prion protein missense variant to Gerstmann-Sträussler syndrome. Nature 338:342–345. doi: 10.1038/338342a0

33. Hsiao KK, Scott M, Foster D, Groth DF, DeArmond SJ, Prusiner SB (1990) Spontaneous neurodegeneration in transgenic mice with mutant prion protein. Science 250:1587–1590. doi: 10.1126/science.1980379

34. Huang AY, Zhang Z, Ye AY, Dou Y, Yan L, Yang X, Zhang Y, Wei L (2017) MosaicHunter: accurate detection of postzygotic single-nucleotide mosaicism through next-generation sequencing of unpaired, trio, and paired samples. Nucleic Acids Res 45:e76. doi: 10.1093/nar/gkx024

35. Jones E, Hummerich H, Viré E, Uphill J, Dimitriadis A, Speedy H, Campbell T, Norsworthy P, Quinn L, Whitfield J, Linehan J, Jaunmuktane Z, Brandner S, Jat P, Nihat A, How Mok T, Ahmed P, Collins S, Stehmann C, Sarros S, Kovacs GG, Geschwind MD, Golubjatnikov A, Frontzek K, Budka H, Aguzzi A, Karamujić-Čomić H, van der Lee SJ, Ibrahim-Verbaas CA, van Duijn CM, Sikorska B, Golanska E, Liberski PP, Calero M, Calero O, Sanchez-Juan P, Salas A, Martinón-Torres F, Bouaziz-Amar E, Haïk S, Laplanche J-L, Brandel J-P, Amouyel P, Lambert J-C, Parchi P, Bartoletti-Stella A, Capellari S, Poleggi A, Ladogana A, Pocchiari M, Aneli S, Matullo G, Knight R, Zafar S, Zerr I, Booth S, Coulthart MB, Jansen GH, Glisic K, Blevins J, Gambetti P, Safar J, Appleby B, Collinge J, Mead S (2020) Identification of novel risk loci and causal insights for sporadic Creutzfeldt-Jakob disease: a genome-wide association study. Lancet Neurol 19:840–848. doi: 10.1016/S1474-4422(20)30273-8

36. Karczewski KJ, Francioli LC, Tiao G, Cummings BB, Alföldi J, Wang Q, Collins RL, Laricchia KM, Ganna A, Birnbaum DP, Gauthier LD, Brand H, Solomonson M, Watts NA, Rhodes D, Singer-Berk M, England EM, Seaby EG, Kosmicki JA, Walters RK, Tashman K, Farjoun Y, Banks E, Poterba T, Wang A, Seed C, Whiffin N, Chong JX, Samocha KE, Pierce-Hoffman E, Zappala Z, O’Donnell-Luria AH, Minikel EV, Weisburd B, Lek M, Ware JS, Vittal C, Armean IM, Bergelson L, Cibulskis K, Connolly KM, Covarrubias M, Donnelly S, Ferriera S, Gabriel S, Gentry J, Gupta N, Jeandet T, Kaplan D, Llanwarne C, Munshi R, Novod S, Petrillo N, Roazen D, Ruano-Rubio V, Saltzman A, Schleicher M, Soto J, Tibbetts K, Tolonen C, Wade G, Talkowski ME, Genome Aggregation Database Consortium, Neale BM, Daly MJ, MacArthur DG (2020) The mutational constraint spectrum quantified from variation in 141,456 humans. Nature 581:434–443. doi: 10.1038/s41586-020-2308-7

37. Keogh MJ, Wei W, Aryaman J, Walker L, van den Ameele J, Coxhead J, Wilson I, Bashton M, Beck J, West J, Chen R, Haudenschild C, Bartha G, Luo S, Morris CM, Jones NS, Attems J, Chinnery PF (2018) High prevalence of focal and multi-focal somatic genetic variants in the human brain. Nat Commun 9:4257. doi: 10.1038/s41467-018-06331-w

38. Khoshkhoo S, Wang Y, Chahine Y, Erson-Omay EZ, Robert SM, Kiziltug E, Damisah EC, Nelson-Williams C, Zhu G, Kong W, Huang AY, Stronge E, Phillips HW, Chhouk BH, Bizzotto S, Chen MH, Adikari TN, Ye Z, Witkowski T, Lai D, Lee N, Lokan J, Scheffer IE, Berkovic SF, Haider S, Hildebrand MS, Yang E, Gunel M, Lifton RP, Richardson RM, Blümcke I, Alexandrescu S, Huttner A, Heinzen EL, Zhu J, Poduri A, DeLanerolle N, Spencer DD, Lee EA, Walsh CA, Kahle KT (2023) Contribution of Somatic Ras/Raf/Mitogen-Activated Protein Kinase Variants in the Hippocampus in Drug-Resistant Mesial Temporal Lobe Epilepsy. JAMA Neurol. 80:578–587

39. Kim, C.C., Haldiman, T., Langeveld, J., Kong, Q.Q., Safar, J.G. (2013) Coexistence and evolution of prions by natural selection. Prion. doi: 10.4161/pri.24863

40. Kim C, Haldiman T, Cohen Y, Chen W, Blevins J, Sy M-S, Cohen M, Safar JG (2011) Protease-sensitive conformers in broad spectrum of distinct PrPSc structures in sporadic Creutzfeldt-Jakob disease are indicator of progression rate. PLoS Pathog 7:e1002242. doi: 10.1371/journal.ppat.1002242

41. Kim C, Haldiman T, Surewicz K, Cohen Y, Chen W, Blevins J, Sy M-S, Cohen M, Kong Q, Telling GC, Surewicz WK, Safar JG (2012) Small protease sensitive oligomers of PrPSc in distinct human prions determine conversion rate of PrP(C). PLoS Pathog 8:e1002835. doi: 10.1371/journal.ppat.1002835

42. Kim C, Xiao X, Chen S, Haldiman T, Smirnovas V, Kofskey D, Warren M, Surewicz K, Maurer NR, Kong Q, Surewicz W, Safar JG (2018) Artificial strain of human prions created in vitro. Nat Commun 9:2166. doi: 10.1038/s41467-018-04584-z

43. Kim J, Park SM, Koh HY, Ko A, Kang H-C, Chang WS, Kim DS, Lee JH (2023) Threshold of somatic mosaicism disrupting the brain function. bioRxiv 2023.12.30.573716

44. Kobayashi A, Teruya K, Matsuura Y, Shirai T, Nakamura Y, Yamada M, Mizusawa H, Mohri S, Kitamoto T (2015) The influence of PRNP polymorphisms on human prion disease susceptibility: an update. Acta Neuropathol 130:159–170. doi: 10.1007/s00401-015-1447-7

45. Kroll F, Dimitriadis A, Campbell T, Darwent L, Collinge J, Mead S, Vire E (2022) Prion protein gene mutation detection using long-read Nanopore sequencing. Sci Rep 12:8284. doi: 10.1038/s41598-022-12130-7

46. Li H, Durbin R (2010) Fast and accurate long-read alignment with Burrows-Wheeler transform. Bioinformatics 26:589–595. doi: 10.1093/bioinformatics/btp698

47. Liu MH, Costa B, Choi U, Bandler RC, Lassen E, Grońska-Pęski M, Schwing A, Murphy ZR, Rosenkjær D, Picciotto S, Bianchi V, Stengs L, Edwards M, Loh CA, Truong TK, Brand RE, Pastinen T, Wagner JR, Skytte A-B, Tabori U, Shoag JE, Evrony GD (2023) Single-strand mismatch and damage patterns revealed by single-molecule DNA sequencing. bioRxiv. doi: 10.1101/2023.02.19.526140

48. Lloyd SE, Mead S, Collinge J (2013) Genetics of prion diseases. Curr Opin Genet Dev 23:345–351. doi: 10.1016/j.gde.2013.02.012

49. Lodato MA, Rodin RE, Bohrson CL, Coulter ME, Barton AR, Kwon M, Sherman MA, Vitzthum CM, Luquette LJ, Yandava CN, Yang P, Chittenden TW, Hatem NE, Ryu SC, Woodworth MB, Park PJ, Walsh CA (2018) Aging and neurodegeneration are associated with increased mutations in single human neurons. Science 359:555–559. doi: 10.1126/science.aao4426

50. Lodato MA, Woodworth MB, Lee S, Evrony GD, Mehta BK, Karger A, Lee S, Chittenden TW, D’Gama AM, Cai X, Luquette LJ, Lee E, Park PJ, Walsh CA (2015) Somatic mutation in single human neurons tracks developmental and transcriptional history. Science 350:94–98. doi: 10.1126/science.aab1785

51. Luquette LJ, Miller MB, Zhou Z, Bohrson CL, Zhao Y, Jin H, Gulhan D, Ganz J, Bizzotto S, Kirkham S, Hochepied T, Libert C, Galor A, Kim J, Lodato MA, Garaycoechea JI, Gawad C, West J, Walsh CA, Park PJ (2022) Single-cell genome sequencing of human neurons identifies somatic point mutation and indel enrichment in regulatory elements. Nat Genet 54:1564–1571. doi: 10.1038/s41588-022-01180-2

52. Martincorena I, Fowler JC, Wabik A, Lawson ARJ, Abascal F, Hall MWJ, Cagan A, Murai K, Mahbubani K, Stratton MR, Fitzgerald RC, Handford PA, Campbell PJ, Saeb-Parsy K, Jones PH (2018) Somatic mutant clones colonize the human esophagus with age. Science 362:911–917. doi: 10.1126/science.aau3879

53. Masters CL, Richardson EP (1978) Subacute Spongiform Encephalopathy (Creutzfeldt-Jakob Disease): The Nature and Progression of Spongiform Change. Brain 101:333–344. doi: 10.1093/brain/101.2.333

54. Mead S, Lloyd S, Collinge J (2019) Genetic Factors in Mammalian Prion Diseases. Annu Rev Genet 53:117–147. doi: 10.1146/annurev-genet-120213-092352

55. Mead S, Poulter M, Beck J, Webb TEF, Campbell TA, Linehan JM, Desbruslais M, Joiner S, Wadsworth JDF, King A, Lantos P, Collinge J (2006) Inherited prion disease with six octapeptide repeat insertional mutation--molecular analysis of phenotypic heterogeneity. Brain 129:2297–2317. doi: 10.1093/brain/awl226

56. Mead S, Poulter M, Uphill J, Beck J, Whitfield J, Webb TEF, Campbell T, Adamson G, Deriziotis P, Tabrizi SJ, Hummerich H, Verzilli C, Alpers MP, Whittaker JC, Collinge J (2009) Genetic risk factors for variant Creutzfeldt-Jakob disease: a genome-wide association study. Lancet Neurol 8:57–66. doi: 10.1016/S1474-4422(08)70265-5

57. Mead S, Uphill J, Beck J, Poulter M, Campbell T, Lowe J, Adamson G, Hummerich H, Klopp N, Rückert I-M, Wichmann H-E, Azazi D, Plagnol V, Pako WH, Whitfield J, Alpers MP, Whittaker J, Balding DJ, Zerr I, Kretzschmar H, Collinge J (2012) Genome-wide association study in multiple human prion diseases suggests genetic risk factors additional to PRNP. Hum Mol Genet 21:1897–1906. doi: 10.1093/hmg/ddr607

58. Mead S, Whitfield J, Poulter M, Shah P, Uphill J, Campbell T, Al-Dujaily H, Hummerich H, Beck J, Mein CA, Verzilli C, Whittaker J, Alpers MP, Collinge J (2009) A novel protective prion protein variant that colocalizes with kuru exposure. N Engl J Med 361:2056–2065. doi: 10.1056/NEJMoa0809716

59. Medori R, Tritschler HJ, LeBlanc A, Villare F, Manetto V, Chen HY, Xue R, Leal S, Montagna P, Cortelli P (1992) Fatal familial insomnia, a prion disease with a mutation at codon 178 of the prion protein gene. N Engl J Med 326:444–449. doi: 10.1056/NEJM199202133260704

60. Meyer M, Kircher M (2010) Illumina sequencing library preparation for highly multiplexed target capture and sequencing. Cold Spring Harb Protoc 2010:db.prot5448. doi: 10.1101/pdb.prot5448

61. Miller MB, Geoghegan JC, Supattapone S (2011) Dissociation of infectivity from seeding ability in prions with alternate docking mechanism. PLoS Pathog 7:e1002128. doi: 10.1371/journal.ppat.1002128

62. Miller MB, Huang AY, Kim J, Zhou Z, Kirkham SL, Maury EA, Ziegenfuss JS, Reed HC, Neil JE, Rento L, Ryu SC, Ma CC, Luquette LJ, Ames HM, Oakley DH, Frosch MP, Hyman BT, Lodato MA, Lee EA, Walsh CA (2022) Somatic genomic changes in single Alzheimer’s disease neurons. Nature 604:714–722. doi: 10.1038/s41586-022-04640-1

63. Minikel EV, Vallabh SM, Lek M, Estrada K, Samocha KE, Sathirapongsasuti JF, McLean CY, Tung JY, Yu LPC, Gambetti P, Blevins J, Zhang S, Cohen Y, Chen W, Yamada M, Hamaguchi T, Sanjo N, Mizusawa H, Nakamura Y, Kitamoto T, Collins SJ, Boyd A, Will RG, Knight R, Ponto C, Zerr I, Kraus TFJ, Eigenbrod S, Giese A, Calero M, de Pedro-Cuesta J, Haïk S, Laplanche J-L, Bouaziz-Amar E, Brandel J-P, Capellari S, Parchi P, Poleggi A, Ladogana A, O’Donnell-Luria AH, Karczewski KJ, Marshall JL, Boehnke M, Laakso M, Mohlke KL, Kähler A, Chambert K, McCarroll S, Sullivan PF, Hultman CM, Purcell SM, Sklar P, van der Lee SJ, Rozemuller A, Jansen C, Hofman A, Kraaij R, van Rooij JGJ, Ikram MA, Uitterlinden AG, van Duijn CM, Exome Aggregation Consortium (ExAC), Daly MJ, MacArthur DG (2016) Quantifying prion disease penetrance using large population control cohorts. Sci Transl Med 8:322ra9. doi: 10.1126/scitranslmed.aad5169

64. Minikel EV, Zhao HT, Le J, O’Moore J, Pitstick R, Graffam S, Carlson GA, Kavanaugh MP, Kriz J, Kim JB, Ma J, Wille H, Aiken J, McKenzie D, Doh-Ura K, Beck M, O’Keefe R, Stathopoulos J, Caron T, Schreiber SL, Carroll JB, Kordasiewicz HB, Cabin DE, Vallabh SM (2020) Prion protein lowering is a disease-modifying therapy across prion disease stages, strains and endpoints. Nucleic Acids Res 48:10615–10631. doi: 10.1093/nar/gkaa616

65. Murley AG, Nie Y, Golder Z, Keogh MJ, Smith C, Ironside JW, Chinnery PF (2023) High-Depth PRNP Sequencing in Brains With Sporadic Creutzfeldt-Jakob Disease. Neurol Genet 9:e200054. doi: 10.1212/NXG.0000000000200054

66. Nichols KE, Malkin D, Garber JE, Fraumeni JF Jr, Li FP (2001) Germ-line p53 mutations predispose to a wide spectrum of early-onset cancers. Cancer Epidemiol Biomarkers Prev 10:83–87

67. Nicolas G, Acuña-Hidalgo R, Keogh MJ, Quenez O, Steehouwer M, Lelieveld S, Rousseau S, Richard A-C, Oud MS, Marguet F, Laquerrière A, Morris CM, Attems J, Smith C, Ansorge O, Al Sarraj S, Frebourg T, Campion D, Hannequin D, Wallon D, Gilissen C, Chinnery PF, Veltman JA, Hoischen A (2018) Somatic variants in autosomal dominant genes are a rare cause of sporadic Alzheimer’s disease. Alzheimers Dement 14:1632–1639. doi: 10.1016/j.jalz.2018.06.3056

68. Oldoni E, Fumagalli GG, Serpente M, Fenoglio C, Scarioni M, Arighi A, Bruno G, Talarico G, Confaloni A, Piscopo P, Nacmias B, Sorbi S, Rainero I, Rubino E, Pinessi L, Binetti G, Ghidoni R, Benussi L, Grande G, Arosio B, Bursey D, Kauwe JS, Cioffi SM, Arcaro M, Mari D, Mariani C, Scarpini E, Galimberti D (2016) PRNP P39L Variant is a Rare Cause of Frontotemporal Dementia in Italian Population. J Alzheimers Dis 50:353–357. doi: 10.3233/JAD-150863

69. Owen F, Poulter M, Shah T, Collinge J, Lofthouse R, Baker H, Ridley R, McVey J, Crow TJ (1990) An in-frame insertion in the prion protein gene in familial Creutzfeldt-Jakob disease. Brain Res Mol Brain Res 7:273–276. doi: 10.1016/0169-328x(90)90038-f

70. Palmer MS, Dryden AJ, Hughes JT, Collinge J (1991) Homozygous prion protein genotype predisposes to sporadic Creutzfeldt-Jakob disease. Nature 352:340–342. doi: 10.1038/352340a0

71. Palmer MS, Mahal SP, Campbell TA, Hill AF, Sidle KC, Laplanche JL, Collinge J (1993) Deletions in the prion protein gene are not associated with CJD. Hum Mol Genet 2:541–544. doi: 10.1093/hmg/2.5.541

72. Parchi P, Castellani R, Capellari S, Ghetti B, Young K, Chen SG, Farlow M, Dickson DW, Sima AA, Trojanowski JQ, Petersen RB, Gambetti P (1996) Molecular basis of phenotypic variability in sporadic Creutzfeldt-Jakob disease. Ann Neurol 39:767–778. doi: 10.1002/ana.410390613

73. Poduri A, Evrony GD, Cai X, Elhosary PC, Beroukhim R, Lehtinen MK, Hills LB, Heinzen EL, Hill A, Hill RS, Barry BJ, Bourgeois BFD, Riviello JJ, Barkovich AJ, Black PM, Ligon KL, Walsh CA (2012) Somatic activation of AKT3 causes hemispheric developmental brain malformations. Neuron 74:41–48. doi: 10.1016/j.neuron.2012.03.010

74. Rivière J-B, Mirzaa GM, O’Roak BJ, Beddaoui M, Alcantara D, Conway RL, St-Onge J, Schwartzentruber JA, Gripp KW, Nikkel SM, Worthylake T, Sullivan CT, Ward TR, Butler HE, Kramer NA, Albrecht B, Armour CM, Armstrong L, Caluseriu O, Cytrynbaum C, Drolet BA, Innes AM, Lauzon JL, Lin AE, Mancini GMS, Meschino WS, Reggin JD, Saggar AK, Lerman-Sagie T, Uyanik G, Weksberg R, Zirn B, Beaulieu CL, Finding of Rare Disease Genes (FORGE) Canada Consortium, Majewski J, Bulman DE, O’Driscoll M, Shendure J, Graham JM Jr, Boycott KM, Dobyns WB (2012) De novo germline and postzygotic mutations in AKT3, PIK3R2 and PIK3CA cause a spectrum of related megalencephaly syndromes. Nat Genet 44:934–940. doi: 10.1038/ng.2331

75. Rohland N, Reich D (2012) Cost-effective, high-throughput DNA sequencing libraries for multiplexed target capture. Genome Res 22:939–946. doi: 10.1101/gr.128124.111

76. Safar JG, Geschwind MD, Deering C, Didorenko S, Sattavat M, Sanchez H, Serban A, Vey M, Baron H, Giles K, Miller BL, Dearmond SJ, Prusiner SB (2005) Diagnosis of human prion disease. Proc Natl Acad Sci U S A 102:3501–3506. doi: 10.1073/pnas.0409651102

77. Safar J, Wille H, Itri V, Groth D, Serban H, Torchia M, Cohen FE, Prusiner SB (1998) Eight prion strains have PrP(Sc) molecules with different conformations. Nat Med 4:1157–1165. doi: 10.1038/2654

78. Sala Frigerio C, Lau P, Troakes C, Deramecourt V, Gele P, Van Loo P, Voet T, De Strooper B (2015) On the identification of low allele frequency mosaic mutations in the brains of Alzheimer’s disease patients. Alzheimers Dement 11:1265–1276. doi: 10.1016/j.jalz.2015.02.007

79. Samocha KE, Robinson EB, Sanders SJ, Stevens C, Sabo A, McGrath LM, Kosmicki JA, Rehnström K, Mallick S, Kirby A, Wall DP, MacArthur DG, Gabriel SB, DePristo M, Purcell SM, Palotie A, Boerwinkle E, Buxbaum JD, Cook EH Jr, Gibbs RA, Schellenberg GD, Sutcliffe JS, Devlin B, Roeder K, Neale BM, Daly MJ (2014) A framework for the interpretation of de novo mutation in human disease. Nat Genet 46:944–950. doi: 10.1038/ng.3050

80. Silveira JR, Raymond GJ, Hughson AG, Race RE, Sim VL, Hayes SF, Caughey B (2005) The most infectious prion protein particles. Nature 437:257–261. doi: 10.1038/nature03989

81. Vogelstein B, Papadopoulos N, Velculescu VE, Zhou S, Diaz LA Jr, Kinzler KW (2013) Cancer genome landscapes. Science 339:1546–1558. doi: 10.1126/science.1235122

82. Wang F, Wang X, Yuan C-G, Ma J (2010) Generating a prion with bacterially expressed recombinant prion protein. Science 327:1132–1135. doi: 10.1126/science.1183748

83. Watson N, Brandel J-P, Green A, Hermann P, Ladogana A, Lindsay T, Mackenzie J, Pocchiari M, Smith C, Zerr I, Pal S (2021) The importance of ongoing international surveillance for Creutzfeldt-Jakob disease. Nat Rev Neurol 17:362–379. doi: 10.1038/s41582-021-00488-7

84. Wei W, Keogh MJ, Aryaman J, Golder Z, Kullar PJ, Wilson I, Talbot K, Turner MR, McKenzie C-A, Troakes C, Attems J, Smith C, Sarraj SA, Morris CM, Ansorge O, Jones NS, Ironside JW, Chinnery PF (2019) Frequency and signature of somatic variants in 1461 human brain exomes. Genet Med 21:904–912. doi: 10.1038/s41436-018-0274-3

85. Werner B, Case J, Williams MJ, Chkhaidze K, Temko D, Fernández-Mateos J, Cresswell GD, Nichol D, Cross W, Spiteri I, Huang W, Tomlinson IPM, Barnes CP, Graham TA, Sottoriva A (2020) Measuring single cell divisions in human tissues from multi-region sequencing data. Nat Commun 11:1035. doi: 10.1038/s41467-020-14844-6

86. Won S-Y, Kim Y-C, Jeong B-H (2023) Elevated E200K Somatic Mutation of the Prion Protein Gene (PRNP) in the Brain Tissues of Patients with Sporadic Creutzfeldt-Jakob Disease (CJD). Int J Mol Sci 24. doi: 10.3390/ijms241914831

87. Zanusso G, Liu D, Ferrari S, Hegyi I, Yin X, Aguzzi A, Hornemann S, Liemann S, Glockshuber R, Manson JC, Brown P, Petersen RB, Gambetti P, Sy MS (1998) Prion protein expression in different species: analysis with a panel of new mAbs. Proc Natl Acad Sci U S A 95:8812–8816. doi: 10.1073/pnas.95.15.8812

88. Zerr I, Ladogana A, Mead S, Hermann P, Forloni G, Appleby BS (2024) Creutzfeldt-Jakob disease and other prion diseases. Nat Rev Dis Primers 10:14. doi: 10.1038/s41572-024-00497-y

89. (2014) Human assembly and gene annotation. In: Ensembl. https://useast.ensembl.org/Homo_sapiens/Info/Annotation. Accessed 23 May 2024

