## Supplementary figures and images for "Neuropathologically-directed profiling of *PRNP* somatic and germline variants in sporadic human prion disease"

### Supplemental Figure 1

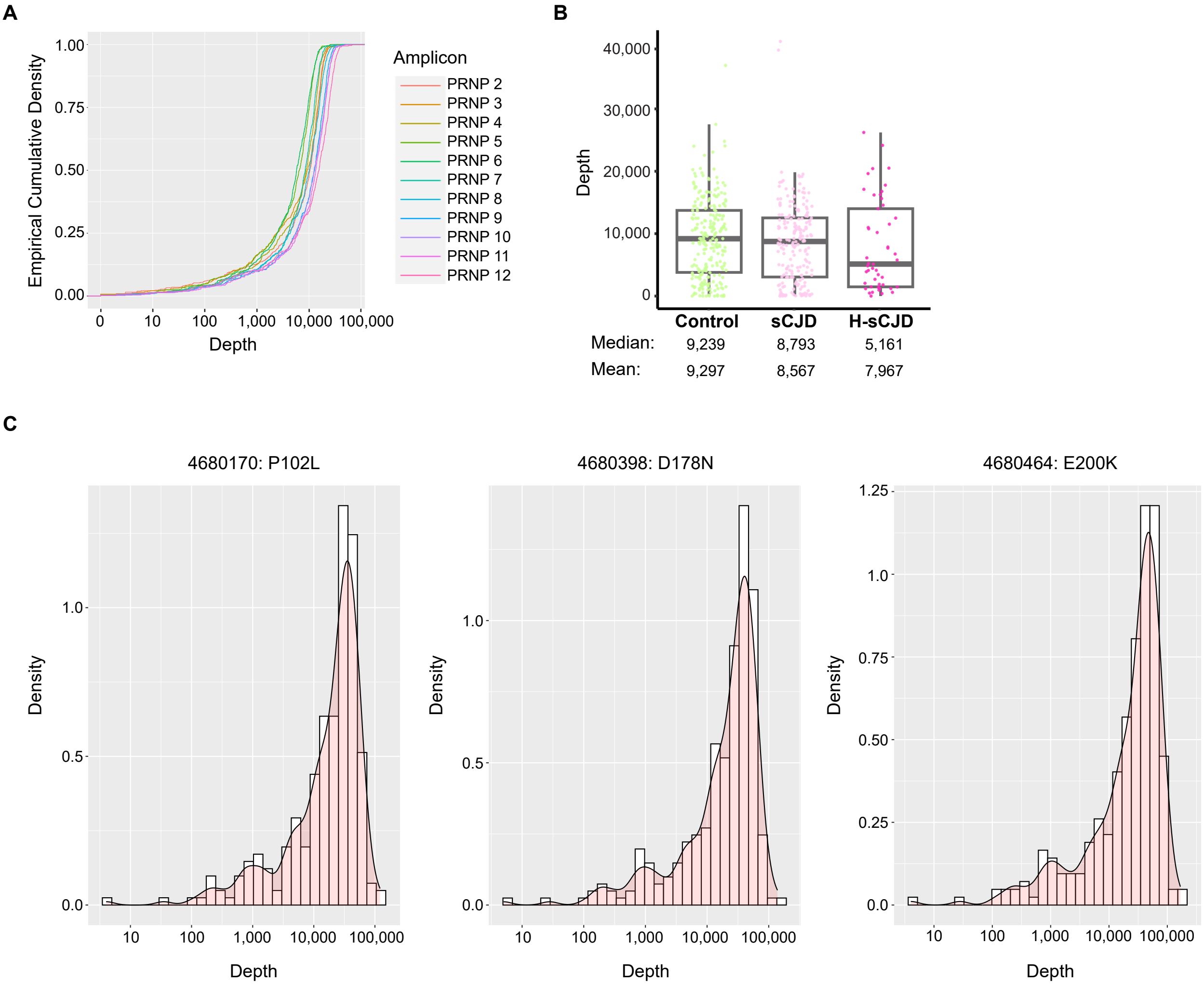

### Supplemental Figure 2

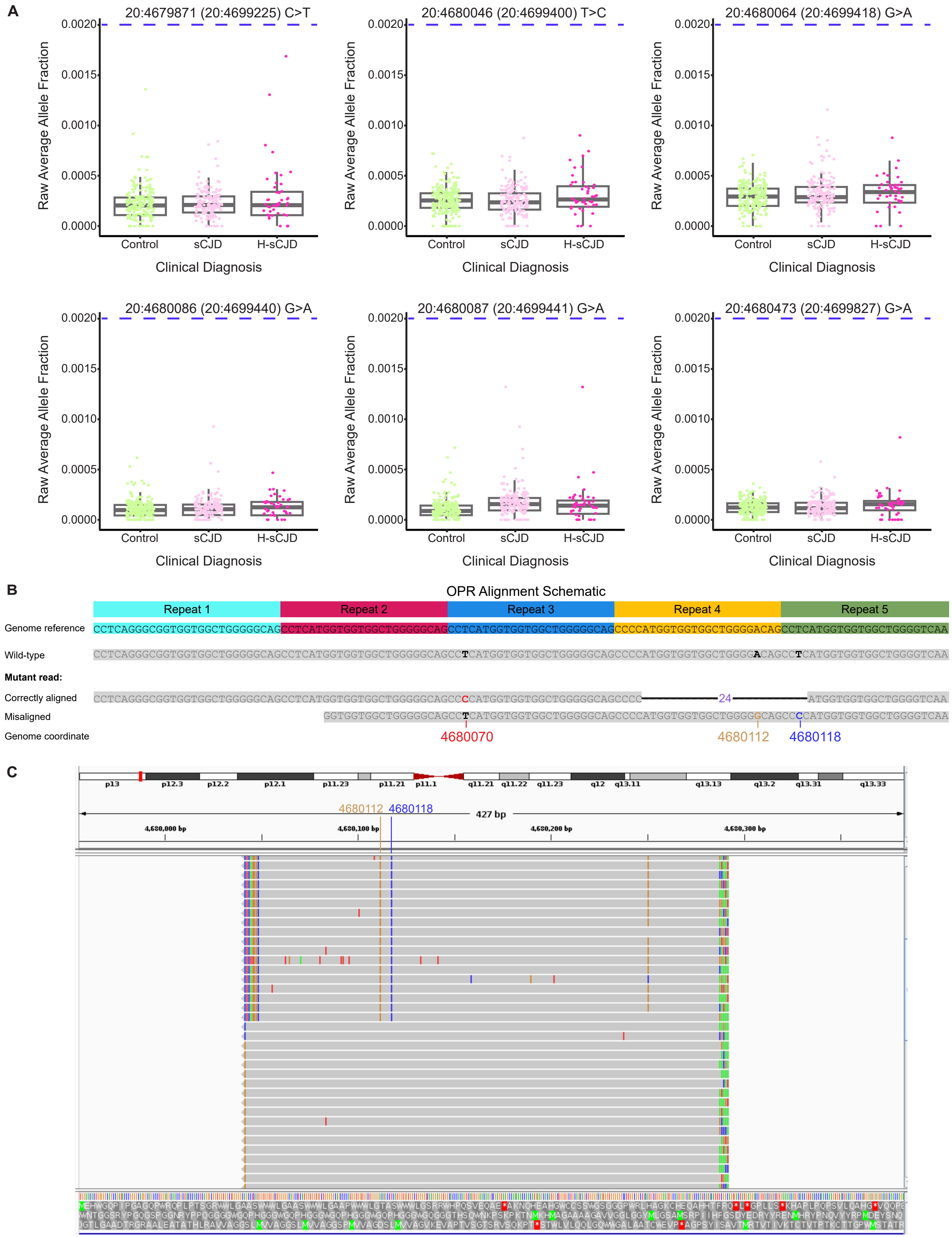

### Supplemental Figure 3

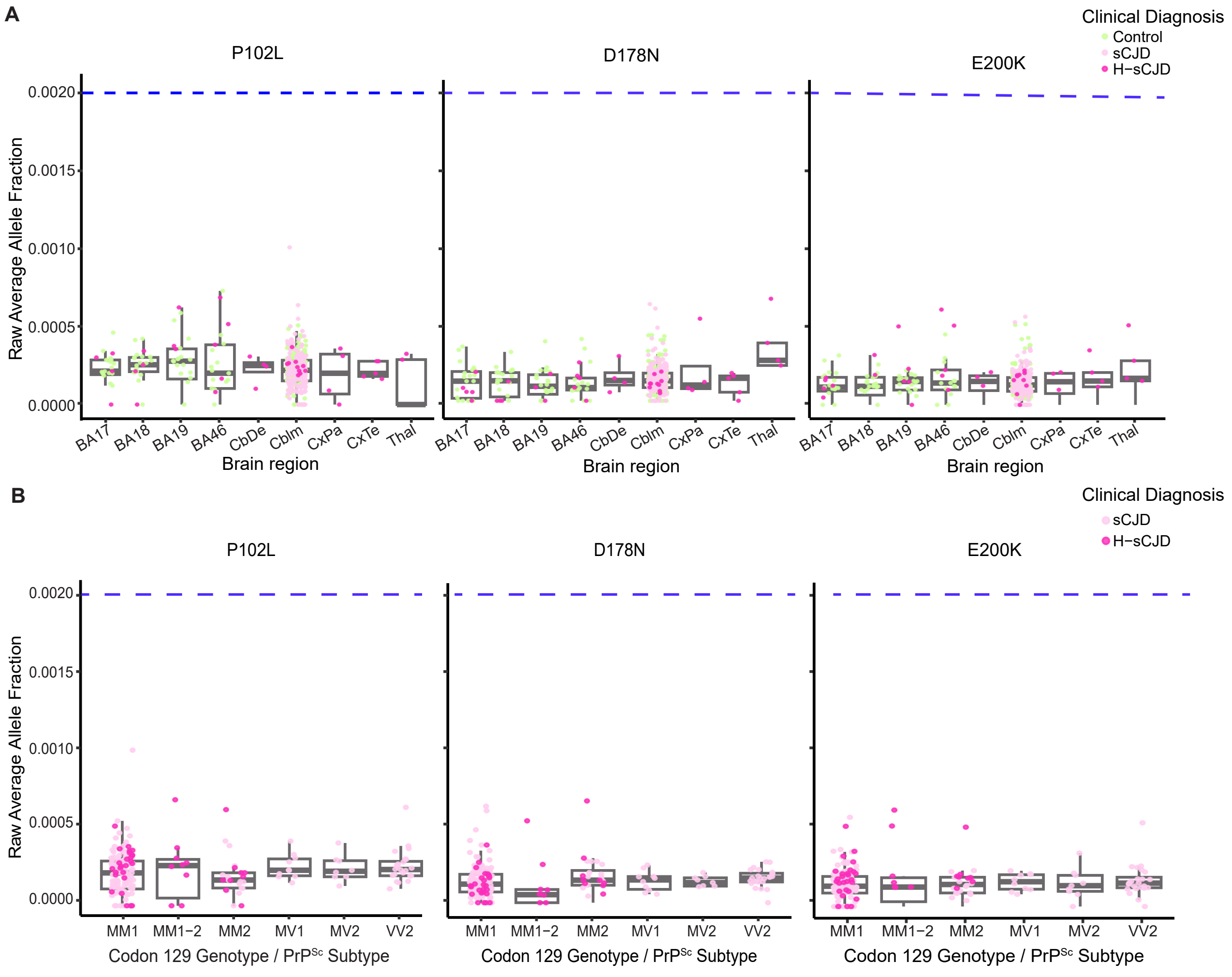
